# Impaired reliability and precision of spiking in adults but not juveniles in a mouse model of Fragile X Syndrome

**DOI:** 10.1101/503714

**Authors:** Deepanjali Dwivedi, Sumantra Chattarji, Upinder S. Bhalla

## Abstract

Fragile X Syndrome (FXS) is the most common source of intellectual disability and autism. Extensive studies have been performed on the network and behavioral correlates of the syndrome but our knowledge about intrinsic conductance changes is still limited. In this study we show a differential effect of FMRP Knock Out (KO) in different sub-sections of hippocampus using whole cell patch clamp in mouse hippocampal slices. We observed no significant change in spike numbers in the CA1 region of hippocampus but a significant increase in CA3, in juvenile mice. However, in adult mice we see a reduction in spike number in the CA1 with no significant difference in CA3. In addition, we see increased variability in spike number in CA1 cells following a variety of steady and modulated current step protocols. This effect emerges in adult (8 weeks) but not juvenile (4 weeks) mice. This increased spiking variability was correlated with reduced spike number and with elevated AHP. The increased AHP arose from elevated SK currents (small conductance calcium activated potassium channels) but other currents involved in mAHP, such as I_h_ and M, were not significantly different. We obtained a partial rescue of the cellular variability phenotype when we blocked SK current using the specific blocker apamin. Our observations provide a single cell correlate of the network observations of response variability and loss of synchronization, and suggest that elevation of SK currents in FXS may provide a partial mechanistic explanation for this difference.

**Significance Statement:** Fragile-X syndrome leads to a range of intellectual disability effects and autism. We have found differential effect of FMRP KO in different sub sections of hippocampus where it caused an increased spiking in CA3 in juveniles and reduced spiking in CA1, in adults. We have also found that even individual neurons with this mutation exhibit increased variability in their activity patterns. Importantly, this effect emerges after six weeks of age in mice. We showed that a specific ion channel protein, SK channel, was partially responsible, and blockage of these channels led to a partial restoration of cellular activity. This is interesting as it provides a possible molecular link between activity variability in single cells, and reported irregularity in network activity.

## INTRODUCTION

Fragile X Syndrome (FXS) is an autism spectrum disorder arising from increased repeats of CGG tri-nucleotide in the FMR1 gene. This leads to silencing of the gene and hence the absence of the FMRP protein. (O’Donnell and Warren, 2002). FMRP is a transcription factor which affects multiple downstream genes including ion channels (Brown et al., 2001; Darnell et al., 2011). Studies have shown that absence of FMRP affects the functioning of ion channels either by regulating the number of channels (Meredith et al., 2007; Strumbos et al., 2010; Gross et al., 2011; Lee et al., 2011; Routh et al., 2013; Ferron et al., 2014; Zhu et al., 2018) or by directly interacting with the channels (Brown et al., 2010; Deng et al., 2013). Multiple studies have shown that potassium channels have a significant effect on spike precision (Fricker and Miles, 2000; Gittelman, 2006; Cudmore et al., 2010), and many of these potassium channels are transcription hits for FMRP. This leads to the hypothesis that FXS may alter the functioning of one or multiple potassium channels, leading to effects on spike precision.

In recent years a number of in-vivo hippocampal recording studies have shown that there is poor correlation of spiking activity between cells, and abnormal theta-gamma phase coupling in FXS mice (Radwan et al., 2016; Arbab et al., 2018; Talbot et al., 2018). In medial Prefrontal Cortex (mPFC), variability in calcium (Ca^2+^) responses has also been observed, leading to impaired Spike Timing Dependent Plasticity (STDP) (Meredith et al., 2007).These studies have led to the ‘discoordination hypothesis’ for FXS (Talbot et al., 2018). This hypothesis states that neurons in FXS are uncorrelated and have aberrant network discharges. In apparent contradiction to this hypothesis, neurons showed hyper connectivity and synchronization in cortical networks of FXS model mice (Testa-Silva et al., 2012; Gonçalves et al., 2013)

Synchronicity is an emergent property of a network and is a function of both network connectivity and intrinsic properties. Specifically, potassium conductances have been shown to have significant effects on spike precision and network synchrony (Fricker and Miles, 2000; Pfeuty et al., 2003; Deister et al., 2009; Cudmore et al., 2010; Gastrein et al., 2011; Hou et al., 2012). Modeling studies have also shown that conductances which mediate spike frequency adaptation help to synchronize network firing (Crook et al., 1998). I_h_ currents, I_M_ currents and SK currents have been shown to mediate adaptation for phase precession in hippocampal neurons (Orban et al., 2006; Kwag et al., 2014), spiking reliability in insect neurons (Gabbiani and Dewell, 2018) and also to maintain regular firing in robust nucleus of acropallium in zebra finches (Hou et al., 2012). Thus FXS mutations, through their effects on potassium channels, may lead to several of the observed network dysfunctions in FXS models.

In the present study, we tested alterations in intrinsic properties of the cell that might underlie the above network observations of uncorrelated activity. First, we observed that cells from FXS knockout (KO) hippocampal slices were more unreliable both at within cell and between cell level, than controls. Consistent with the developmental manifestations of FXS, this effect was absent in juvenile mice and emerged only after ∼6 weeks. Decreased spiking in KO mice was correlated with increased variability. At the mechanistic level, we found that the spiking changes were due to elevated medium AHP (mAHP) which in turn was due to elevated SK currents. In contrast to these observations of decreased spiking in the CA1 region of adult animals, we observed increased spiking in the CA3 region of juvenile animals (as seen in Deng et al., 2019) but not in CA1. We did not see any significant change in spatial distribution of SK channels either in CA3 or CA1, in either juvenile animals or in adult animals. This too is consistent with earlier observations (Deng et al., 2019). Finally, we tested the outcome of blockers of SK and found that that they indeed led to a partial rescue of the observed phenotype of increased spiking variability in adult animals.

## Materials and Methods

### Animals

C57BL/6 strain of *Fmr1tmCgr* male mice were used for the experiments. All experimental procedures were approved by the NCBS ethics committee (Project ID: NCBS-IAE-2017/04(N)). The animals were housed in the institute animal house where they were maintained in 12 hr light and 12 hr dark cycle. The animals used older animal group was in the range of 6-8 weeks of age, and the younger group was 3-4 weeks old.

### Slice preparation

Mice were anaesthetized with halothane. Their head was decapitated after they were euthanized by cervical dislocation. Hippocampal slices were made in the ice cold aCSF of the composition: 115 mM NaCl, 25 mM glucose, 25.5 mM NaHCO_3_, 1.05mM NaH_2_PO_4_, 3.3mM KCl, 2mM CaCl_2_, and 1mM MgCl_2_. 400μm thick slices were made using a VT1200S vibratome and then incubated at room temperature for 1 hour in the aCSF, which was constantly bubbled with 95% O_2_ and 5%CO_2_. Subsequently, the slices were transferred to the recording chamber where they were maintained at an elevated temperature of 30⁰C-34⁰C for the recordings.

### Electrophysiology

CA1 neurons were identified under an upright DIC microscope (Olympus BX1WI microscope) using a 40X objective (water immersion lens, 0.9 NA LUMPLFLN, 40X). 2-4 MOhm pipettes were pulled from thick walled borosilicate glass capillaries on a P-1000 Flaming micropipette puller (Sutter instruments). The pipettes were filled with internal solution of the following composition for whole cell Current Clamp recordings: 120mM potassium gluconate, 20mM KCl, 0.2mM EGTA, 4mM NaCl,10mM HEPES buffer, 10mM phosphocreatine, 4mM Mg-ATP, and 0.3mM Na-GTP (pH 7.4, 295 mOsm). For Voltage Clamp recordings, the same composition of internal solution was used with the one change that 120mM potassium gluconate was substituted with 120mM potassium methylsulphate. Cells were recorded if they had a resting potential <-60mV. We also required that they exhibit stable firing with little or no depolarization block for lower current inputs. Series resistance and input resistance were continuously monitored during the protocols and the cell was discarded if these parameters changed by >25%.

### Protocol for measuring spike variability and analysis

All spike variability and precision experiments were done in current clamp mode. A step input current stimulus of 150pA DC for 900 ms was used for the majority of the recordings. In some cases, as indicated in the text, frozen noise and sinusoidal input currents were also used, riding on a baseline current step of 150pA, and again for a duration of 900 ms. SD of noise used in noise protocols were 10pA, 25pA, 50pA and 100pA with a time-cutoff of τ=3msec. For sinusoidal currents SD of 50pA and 100pA were used at 5 Hz. All the protocols were repeated for 25 trials. Within-cell spike variability (CV_w_) was computed as Coefficient of Variation (CV) in spike numbers across 25 trials. Spike numbers computed for all 25 trials were used to find the Standard Deviation in spike numbers which was then divided by mean number of spikes to find CV_w_.

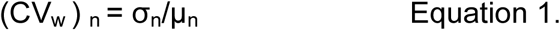

Where *n* is the cell index, σ*_n_* is the standard deviation across trials within cell *n*, and µ*_n_* is the mean number of spikes across all the trials within cell *n*.

The vector CV_w_ was computed for all recorded KO and WT cells respectively, and used for comparing the responses (Fig 1E). The distributions in CV_w_ were compared using the Wilcoxon test.

**Figure 1.**
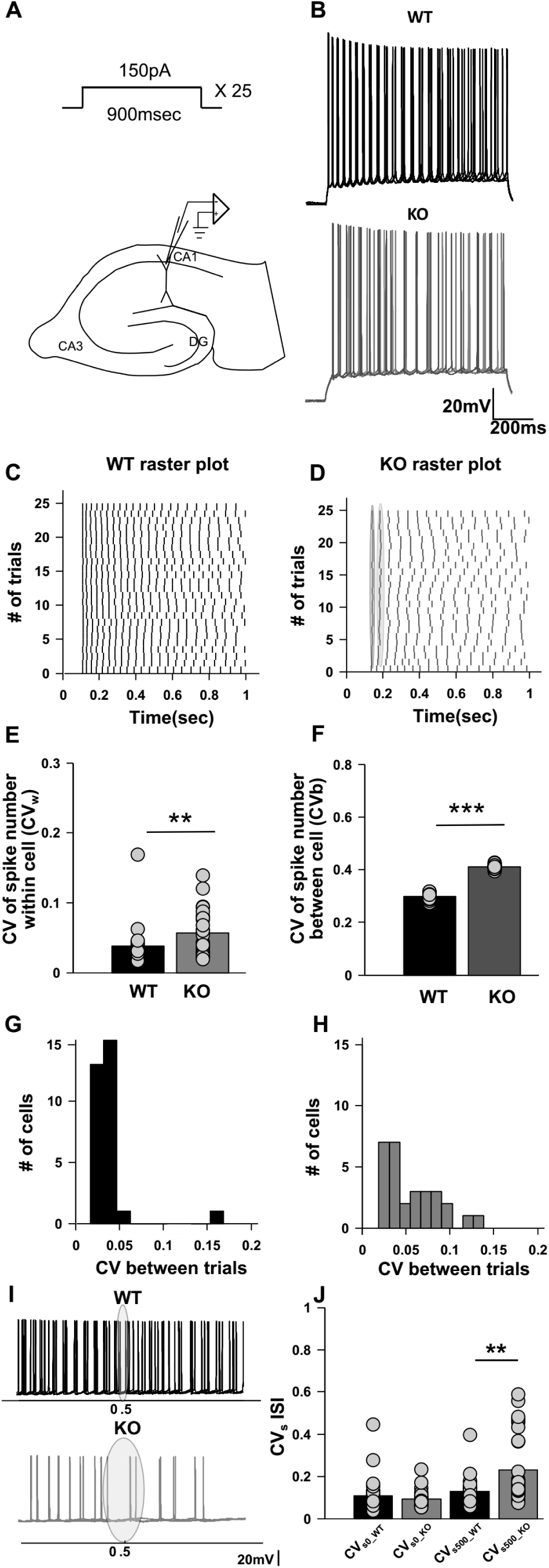
Increased spike variability and spike timing imprecision both at within cell level (CV_w_) across multiple trials and between cells (CV_b_) in CA1 hippocampus of FXS animals. (A) Schematic of a hippocampal slice and the step depolarization protocol. The protocol consists of a 150pA step for 900msec time duration repeated over 25 trials. (B) Representative raw traces of spiking, superimposed for four trials only for WT (black) and KO (grey). Scale bar 20mV, 200msec. (C) Representative raster plot showing spiking for the step depolarization protocol for WT (black) and (D) KO (grey). Also showing the methodology for computing jitter slope for estimating spike timing precision, further plotted in 1I and 1J. (E) Bar plot for within cell variability across multiple trials (CV_w_) WT (n=30 cells, black) and KO (n=29 cells, grey). KO cells have significantly higher within cell variability than WT cells (p=0.0048, Wilcoxon rank sum test). (F) Bar plot for between cell variability (CV_b_) for matched trials, WT (n=30 cells, black) and KO (n=29 cells, grey). KO cells have significantly higher between cell variability than WT cells (p<0.001, Wilcoxon rank sum test) (G) Histogram showing spread of CV_w_ parameter for WT (n=30 cells, black) and (H) for KO (n=29 cells, grey). KO cells have more spread in CV_w_ parameter, depicting heterogeneity in KO cell population. (I) Representative trace showing the time window where CV_s_ in ISI is calculated for WT (upper black trace) and for KO (lower grey trace). Superimposed traces are for four trials only. Scale bar 20mV. (J) Bar plot for CV_s_ in ISI, a measure of spike precision for WT (n=30 cells, black) and KO (n=29 cells, grey). CV_s0_ is not significantly different between WT and KO. However, there is a significant increase in the CV_s500_ for KO cells as compared to WT (p=0.018 for WT vs KO differences; two-way ANOVA

The analysis of CV_b_ was derived from a paper which reports experiments on network synchrony (Talbot et al., 2018). In contrast to our CV_w_ analysis where variability in spiking was assessed within each cell, in CV_b_ we measure across trials between cells. For computing between-cell spiking variability (CV_b_), spike numbers on corresponding trial numbers were compared between different cells. Specifically, we constructed a vector for each cell whose entries were the number of spikes for successive trials. These vectors were compared across different cells, within the same genotype and used to find CV_b_. The following formula was used for the computation:

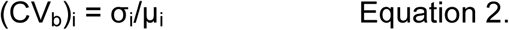

Where *i* is the trial index, σ_i_ is the standard deviation over trial *i* between cells, and µ*_i_* is the mean number of spikes on trial *i*, between all cells. The CV_b_ vectors were computed for WT and KO populations respectively (Figure 1F) and compared using Wilcoxon statistics.

### Measurement of spike timing precision: CV_s_

Spike times were found for all the spikes in all trials. The Inter Spike Interval (ISI) was computed for the first two spikes and for spikes that bracketed a time of 500ms after the start of the step input protocol. For each cell *m* we obtained the Coefficient of variation, CV_s0_m_ and CV_s500_m_ for the first ISI and the ISI at 500 ms, respectively:

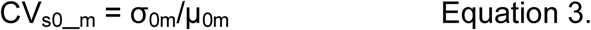

Where *m* is the cell index, σ*_m_* is the standard deviation of ISI across trials within cell *m*, and µ*_m_* is the mean ISI across all the trials for cell *m*. We calculated CV_s500_m_ in a similar manner.

By repeating this calculation over all cells, we obtained the following four vectors: CV_s0_WT_, CV_s500_WT_, CV_s0_KO_, and CV_s500_KO_. The distributions in these four vectors were compared using a two-way ANOVA.

### Measurement of spike timing precision: Jitter Slope

The jitter slope analysis was taken from (Bacci and Huguenard, 2006). Jitter was defined as σ_ti_, which is the standard deviation of spike timing for spike number *i*, compared over all trials for a given cell. In this manner the vector σ_t_ was obtained by considering all the spike numbers. Note that not all trials had the same number of spikes, so the final few spike numbers may have had fewer samples in σ_t_.(Extended figure 1-1).

The Jitter slope was obtained as a regression fit for the vector σ_t_, with the x axis defined by the corresponding vector for mean timings µ_t_. This slope was calculated for spikes falling in the last 400 ms of the current pulse since this is where the SK channel is activated. As a pre hoc criterion, cells were required to have a minimum of two spike time points in order to be selected for this calculation. Note that this criterion excluded several cells which had strong suppression of firing towards the end of the trial, presumably due to SK channel activation. Thus this measure was compromised by being insensitive to precisely those cells with the largest effects in the KO.

(Extended data 1-1)

### Measurement of fI curves

The measurement of f-I curves was done in current clamp. For plotting f-I curves, each cell was given input currents from −100pA to 400pA at its resting membrane potential. The mean ± SEM of the number of spikes elicited for each input current was plotted on the Y axis, and the corresponding input current on the X axis.

### AHP measurement

AHPs were measured by holding the cell at −70mV in Current Clamp mode, using some holding current. In each trial, the holding current was set to zero for 200msec and then step input for 100msec was delivered. The current during this step input was adjusted so that the cell produced 5 spikes in 100msec. This stimulus was repeated for 15 trials and the voltage traces of all the trials were averaged. AHPs were analyzed as the hyperpolarization after the spike train. mAHP was computed as the average hyperpolarization 50msec after the spike train, and sAHP was computed 200msec after the spike train (Fig 5A and 5B).

### Immunofluorescense experiments and analysis

3-4 week or 6-8 week old animals were anaesthetized and brains were dissected out. Brains were fixed in 4%PFA for 48hrs at 4°C. Post fixation, brains were washed with PBS and 30 µm thick slices were cut on a VT12000S vibratome. Slices were kept in cryoprotectant solution and stored at −20°C for further use. Slices were re-suspended in Phosphate Buffer Saline (PBS) solution for the experiment. 50mM Ammonium Chloride (NH_4_Cl) was used to quench auto fluorescence from the samples. After multiple steps of washing and permeabilization with 0.3% PBS with Triton-X 100, cells were incubated with 5% Normal Goat Serum (NGS) for blocking for 1-2 hr. Followed by washes, slices were incubated in primary antibody for SK2 channel, overnight at 4°C. Primary antibody was purchased from Alomone Labs (Cat # APC–028) and were used in 1:300 dilution with 2.5% NGS. Further, slices were incubated in secondary antibody at 1:500 dilution for 1-2 hrs at room temperature. Followed by several washes and DAPI stain (Sigma) for 15 mins in PBS, slices were mounted in Prolong Gold mounting media and imaged in Olympus FV3000 confocal microscope.

### Measurement of M and I_h_ currents

For measuring Ih sag produced in voltage change, cells were held at −70mV and given a step hyperpolarization of 250pA for 500msec, in current clamp mode. Voltage deflection was calculated by the finding the percentage change between maximum and steady state voltages during the hyperpolarizing pulse ((V_max_-V_ss_)/V_max_)/100. Each cell was presented with 15 trials and I_h_ sag was averaged across all the trials to get a value for a cell.

TTX (0.5 µM) was added in the bath while measuring M currents in voltage-clamp. For measuring M currents, cells were held at −20mV and M currents were deactivated in steps from −20mV to −80mV in difference of 10mV of 900msec time duration. Difference between steady state current after the start of the protocol (from the last 250msec) and baseline of the cell was quantified as the current elicited due to the protocol. Specific M current blocker XE991 (30µM) was perfused to get post blocker trace. Difference between Pre and Post XE991 was quantified as M current.

### Measurement of SK currents

TTX (0.5 µM) was added in the bath while measuring SK currents in voltage-clamp. For measuring SK currents cells were held at −55mV while increasing voltage steps were given from −55mV to +35mV, in increments of 10mV, for 800msec duration. Each voltage step was preceded with a small hyperpolarization from −55mV to - 65mV for 100msec. Cell capacitance compensation was done during the recordings. The SK currents were estimated from the amplitude of the outward current 25 ms after the voltage was returned to the holding potential as the initial part of the trace will be contaminated with capacitance currents (Figure 6A and 6B). Apamin perfusion (100nM) was done to identify these currents. Same protocol was used post apamin perfusion. The difference current of pre apamin and post apamin was quantified to find apamin sensitive SK current.

### Data acquisition and Analysis

Recordings were done using an Axopatch 200 B amplifier and digitized using Digidata 1400A. Current clamp recordings were filtered at 10 KHz and sampled at 20 KHz. Voltage Clamp recordings were carried out at a gain of 5, filtered at 3 KHz, sampled at 20 KHz. All analysis was done using custom written code in MATLAB (R2013a) and Graph Pad Prism 6. Statistical tests were done after checking the normality of the data. We plotted the data to check for normality. Even if the data looked normal, Wilcoxon rank sum test was used for most analysis for more stringent results. For multiple comparisons ANOVA was used with Sidak’s post hoc test or t-test with Bonferroni correction as post hoc tests. All the data is plotted as mean ± SEM.

### Drugs and Chemicals used

All toxins were purchased from Sigma other than TTX which was from Hello Bio. TTX (Stock of 0.5mM) and apamin (Stock solution of 1mM) were made in milliQ water. XE991 (Stock 10mM) was made in DMSO. All stocks were stored at −20° C and were diluted to the working concentration immediately before the experiments.

### Experiment design and statistical analysis

We have used hippocampal slice preparation from male mice in this study, where CA1 and CA3 regions were used for electrical recordings. Different current inputs were injected into the cell soma via recording electrode and elicited spikes were recorded in current clamp mode. Spiking variability was assessed by finding Coefficient of Variation (CV) for within cell and between cells. Spike precision was assessed by finding slope of Standard deviation in spike times over multiple trials. SK currents were measured in voltage clamp and were found to be partially responsible for increased variability. All analysis and plotting was done in MATLAB and GraphPad 6. Normality of the data was checked by plotting it. Even if the distribution followed a normal distribution profile, Wilcoxon test was used. For multiple comparisons 2- way or 3- way ANOVA with Sidak’s post hoc test or t-test using Bonferroni correction were used as post hoc test.

## RESULTS

### FMR1 KO cells exhibit increased variability in spike numbers and impaired spike precision

We performed whole cell patch clamp recordings in brain slices from CA1 cells of dorsal hippocampus in male wild-type (WT) and KO littermate animals (6-8weeks old). Somatic current injection was used to inject various waveforms of current (Fig 1A). These included a square current step, frozen noise riding on a current step, and a sinusoidal waveform riding on a current step (Fig 2). In each case we measured the variability in spiking over the 900 ms of current injection, in two respects: first, the variability within a neuron, over repeated trials (CV_w_), and second, the variability between neurons, comparing matching trials between cells (CV_b_) (methods).

**Figure 2.**
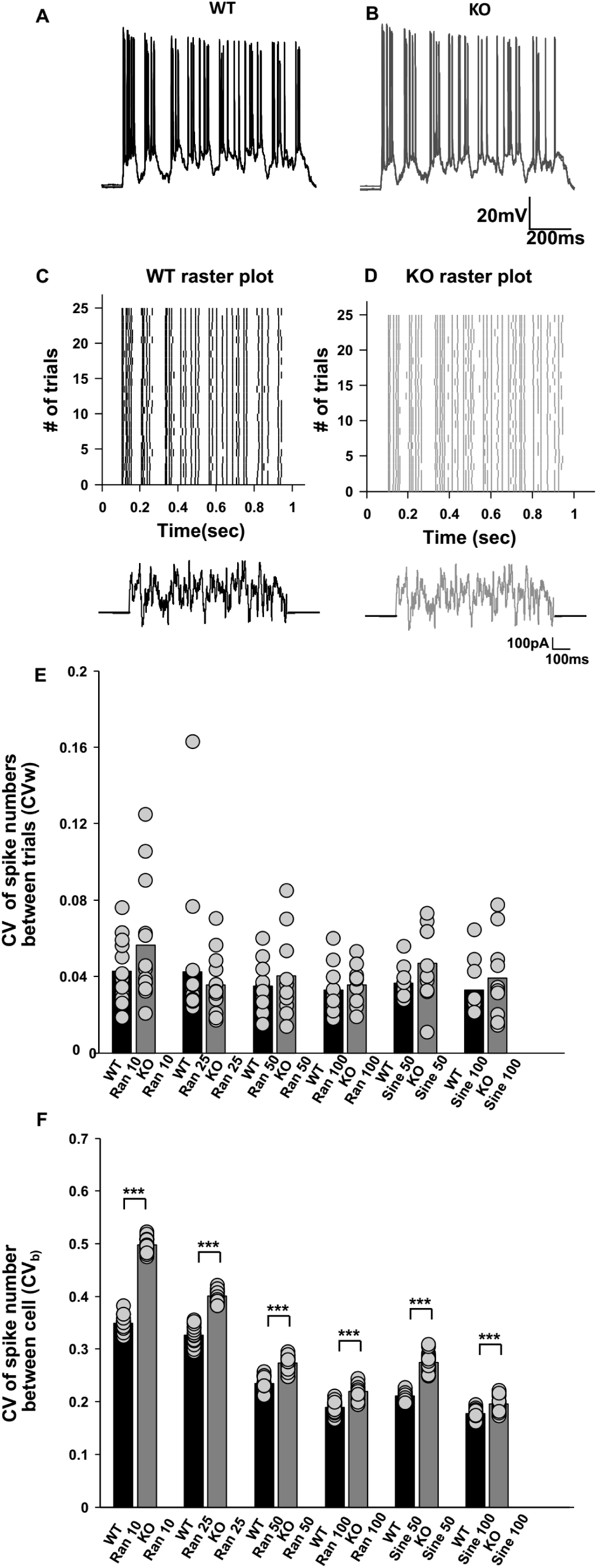
KO cells have increased spike variability between cells (CV_b_) for noise and sinusoidal stimuli but not for within cell variability (CV_w_) across trials. (A) Representative raw plot of spiking showing no spike variability or spike precision differences for Ran 100 noise protocol across multiple trials for WT(black) and (B) for KO (grey). The trace is for four superimposed trials. Scale bar 20mV, 200msec. (C) Raster plot showing no spike variability or spike precision differences for Random 100 noise protocol across multiple trials for WT(black) and (D) for KO (grey). Trace below indicates Ran 100 input protocol. Scale bar 100pA, 100msec (D) Bar plot for within cell variability differences (CV_w_) for all noise and sinusoidal protocols for WT (black bars) and KO (grey bars). The protocol is numbered by the standard deviation (SD) in pA (random protocols) or sine-wave amplitude (sine protocols) (Random 10 protocol: n=16 for WT and n=13 for KO; Random 25: n=16 for WT and n=13 for KO; Random 50: n=10 for both WT and KO; Random 100: n=10 for both WT and KO; Sine 50: n=10 for both WT and KO; Sine 100: n=10 for both WT and KO; error bar represent SEM). No significant within cell variability (CV_w_) differences were seen between WT and KO in any protocol. (E) Bar plot for between cell variability differences (CV_b_) for all noise and sinusoidal protocols (Random 10: n=16 for WT and n=13 for KO; Random 25: n=16 for WT and n=13 for KO; Random 50: n=10 for both WT and KO; Random 100: n=10 for both WT and KO; Sine 50: n=10 for both WT and KO; Sine 100: n=10 for both WT and KO). Significant differences were seen (p < 0.001; three-way ANOVA, error bar represent SEM) for all pairs.

We computed the co-efficient of variation (CV_w_) of spike numbers for each of the current stimulus protocols (methods). We observed that the CV of spike numbers across trials within cells (CV_w_) was significantly higher for KO cells as compared to WT, but only for the step current protocol (CV_w_ for WT 0.038 ± 0.005, n=30 cells; CV_w_ for KO 0.057 ± 0.006, n=29 cells; p=0.0048, Wilcoxon rank sum test) (Fig 1B, 1C,1D,1E). For other protocols, where frozen current noise or sinusoidal waveforms were riding on top of a current pulse, spikes were very reliable over different trials for both WTs and KOs (Fig 2A, Fig 2B, Fig 2C, Fig 2D and Fig 2E). We then estimated the variability between cells (CV_b_). To do this, we compared spike numbers between different cells of WT and KO by matching corresponding trials, such that we compared spiking between the first trials of WT between all the WT cells, second trials of WT between all cells and so on forth, within the same genotype. Similar comparisons were made for KO cells as well (methods). From this procedure we found that between cell variability was significantly elevated in KO cells as compared to WT (CV_b_ for WT cells 0.29 ± 0.003, n=30 cells; CV_b_ for KO cells 0.41 ± 0.002, n=29 cells; p<0.001; Wilcoxon rank sum test) (Fig 1F).We plotted the CV_w_ of spike numbers within the cell, for WT and KO population and observed that KOs have much wider spread as compared to WT (SD of the CV_w_ population for WT cells = 0.026, n=30 cells; SD of the CV_w_ population for KO cells = 0.03, n=29 cells) (Fig 1G and 1H). This implied that KOs have a more heterogeneous population than WT cells. In KO mice, some cells were reliable, i.e., they produced a similar number of spikes across multiple trials but some did not. Based on this analysis (Fig 1G and 1H) we found that 51.7% of KO cells had CV_w_ values which did not overlap with those of WT cells.

Spike timing precision was computed by finding CV_s_ in inter-spike intervals (ISIs, methods). We compared CV_s0_ with CV_s500_ to assess the contribution of SK channels, which are shut at t=0, but open by t=500. We chose not to use the last ISI for this comparison because the timing of the last spike was very variable in KO cells. We then compared CV_s_ at t=0 and t=500, between WT and KO cells using 2-way ANOVA. We found that there was no significant difference in between WT and KO for CV_s0_. However, at t=500, we found that CV_s500_KO_ was significantly elevated compared to CV_s500_WT_ (CV_s0_WT_ = 0.11 ± 0.014; CV_s0_KO_ = 0.09 ± 0.007; CV_s500_WT_ = 0.13 ± 0.01; CV_s500_KO_ = 0.23 ± 0.03; F _(1,113)_ =5.78; p=0.018 for WT vs KO differences; two-way ANOVA) (Fig 1J) Increased within cell (CV_w_), between cell (CV_b_) variability of spike numbers and increased CV_s_ of spike timings are both indicative of abnormal coding in KO neurons. Thus these initial recordings showed that there was increased heterogeneity in spiking in CA1 pyramidal neurons of KO mice.

### KO mice exhibit increased variability in spike number between cells (CV_b_) for frozen noise stimuli

Neuronal coding takes place against a noisy background of continuous synaptic input (Faisal et al., 2008). Frozen noise stimuli have therefore been used in several studies to examine single-neuron computations and are considered more physiological (Stevens and Zador, 1998; Carandini, 2004; Fujisawa et al., 2004; Han and Boyden, 2007). We injected noise waveforms in the soma using a design similar to that of Mainen and Sejnowski, 1995. The stimulus waveforms were mainly either of random noise or sinuosoidal input currents (for details see methods). The resting Vm was controlled by maintaining constant current injection into the cell while recording. Cells were only accepted if the fluctuations in their potential while recording were smaller than ± 2.5mV. For noisy inputs, spikes across multiple trials aligned very precisely for both WT (Fig 2A, 2C) and KO cells (Fig 2B, 2D). Upon computation of CV of spike numbers across trials within cell (CV_w_), there were no statistically significant differences for any of the protocols (Fig 2E). However, between cell variability differences (CV_b_) were still significantly elevated for KO cells, for all stimulus protocols (Fig 2F). We did three way ANOVA for all the protocols on the no of spikes produced in respective protocols for WT and KO. The 3 factors were: across trials, across cells and WT vs. KO. ANOVA results were significant for across cells (p<0.001), for WT vs. KO (p<0.001) but not for across trials (p=1) for all the protocols. Thus, between cell variability (CV_b_) reveals robust phenotypic differences which are also observed between WT and KO for the input noise stimulus. The same is not true for across trials within cell (CV_w_) differences.

### Spike imprecision diverges between KO and WT only in adult mice

Fragile X is known to have an age dependent effect where some disease phenotypes get worse with age (Larson, 2005; Cornish et al., 2008; Choi et al., 2010). We asked if similar effects of age might manifest in the phenotype of firing imprecision of KO cells. We therefore repeated our experimental protocols in young (3-4 week old) male WT and KO littermate mice. Regular spike firing was observed in both KO and WT cells for step input current injection (Fig 3A and 3B). As before, we assessed variability of spike numbers across multiple trials for within the cell (CV_w_).There was no significant difference between WT vs. KO in these younger mice (CV_w_ for WT 0.041 ± 0.006, n=26 cells; CV_w_ for KO 0.03 ± 0.002,n=19 cells; p=0.81; Wilcoxon rank sum test) (Fig 3A,3B,3C,3D and 3E). Spike precision was assessed as above, by finding CV_s_ of ISI. By this measure too, no significant difference in precision of spike timings was found for KO as compared to WT (CV_s500_ for ISI for WT younger 0.133 ± 0.02, n=26 cells; CV_s500_ for ISI for KO younger 0.103 ± 0.01, n=19 cells; p=0.82, Wilcoxon rank sum test) (Fig 3F). We used a two way ANOVA interaction model for assessing age-dependent differences in spiking variability across different trials in a cell (CV_w_) (F _(1, 100)_ = 4.97, p=0.028 for WT and KO differences; F_(1, 100)_ = 8.05, p=0.005 for interaction factor; two-way ANOVA with interaction model) (Fig 3E). We also estimated differences in spike timing precision across different trials in a cell (CV_s500_) (F _(1, 99)_ = 7.47, p=0.007 for age factor, F _(1, 99)_ = 9.46, p=0.003 for interaction factor; two-way ANOVA with interaction model) (Fig 3F). ANOVA indicated a significant interaction between the two factors: age and genotype of the animal. Using post hoc tests we concluded that for KO cells there is a worsening of phenotype for both variability in spiking number (CV_w_) across multiple trials within a cell and in spike timing precision (CV_s500_ of ISI) across multiple trials within a cell (Fig 3E and 3F, detailed statistics shown in table 1). However, the same was not seen in WT cells. A two-way ANOVA was done on the number of spikes produced in the 900msec stimulus period used in the step depolarization protocol (Fig1A). In this ANOVA the two factors were: young vs old animals and WT vs. KO. There was no significant change between ages of the animals for both WT and KO (F _(1, 101)_ = 0.21; p=0.6) but there was a significant reduction in spike number for KO animals as compared to WT (F _(1, 101)_ = 5.5; p=0.021) (Fig 3G). However, the frequency-vs.-current (f-I) curves were not significantly different between WT and KO either in the older (8 weeks) or younger (4 weeks) age groups, when slope of the f-I curves (gain) was used as a readout (Mean gain for f-I curve older WT 0.16 ± 0.007 spikes. s^−1^. pA ^−1^, n= 27 cells; mean gain for f-I curve older KO 0.16 ± 0.008 spikes. s^−1^. pA ^−1^, n= 25 cells; Mean gain for f-I curve younger WT 0.133 ± 0.01 spikes. s^−1^. pA ^−1^, n=26 cells; mean gain for f-I curve younger KO 0.136 ± 0.013 spikes. s^−1^. pA ^−1^, n=19 cells; F _(1, 94)_ = 0.03; p=0.86 for WT and KO differences, two way ANOVA) (Fig 3I and 3J).When comparisons were made of f-I curves of WT and KOs cells, there was reduced excitability seen for some input currents in older animals (Mean # of spikes for input current 125 pA in WT animals 22.07 ± 1.74; n=27 cells; Mean # of spikes for input current 125 pA in KO animals 15.92 ± 2.1; n=25 cells; F _(1, 1070)_ = 31.03, p<0.001; two *way* ANOVA) (Fig 3L), but not in younger animals (Mean # of spikes for input current 125 pA in WT animals 21.8 ± 2.9; n=26 cells; Mean # of spikes for input current 125 pA in KO animals 22.8 ± 2.9; n=19 cells, F _(1, 923)_ = 0.92; p=0.33; two *way* ANOVA) (Fig 3K). Other properties like spike threshold were not significantly different between WT and KO (Spike threshold for the 1^st^ spike WT −53.6 mV ± 0.56 n=30 cells; spike threshold for 1^st^ spike KO −53.8 mV ± 0.62, n=29 cells; Spike threshold for the last spike WT −41.9 mV ± 0.66, n=30 cells; Spike threshold for the last spike KO −43.5 mV ± 0.66, n=29 cells, F _(1, 115)_ = 1.98; p=0.16; two way ANOVA). The spread in number of spikes produced by KO neurons was greater than observed in WT, further substantiating the point of increased variability in KO neurons (SD of spike numbers WT = 7.2; n=750 spikes from 30 cells 25 trials each; SD of spike numbers KO = 8.1; n=725 spikes from 29 cells 25 trials each)(Fig 3H).

**Figure 3.**
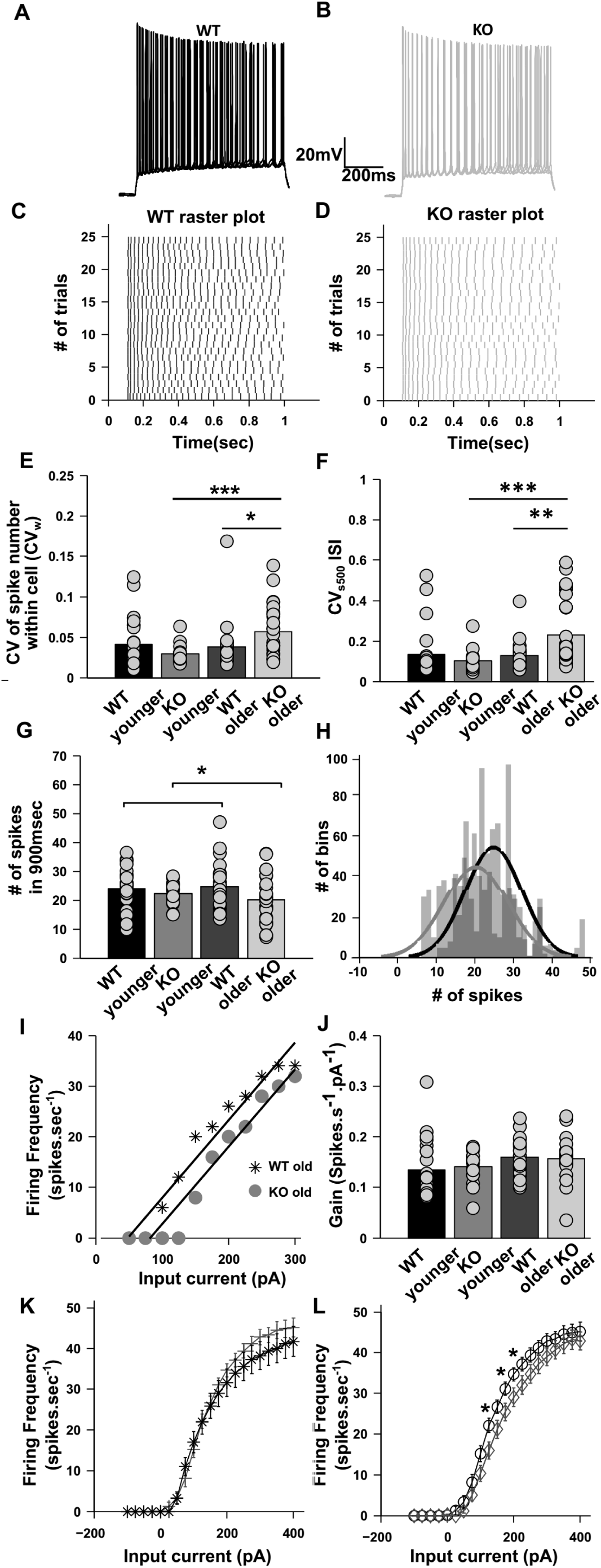
Spiking variability and spike imprecision diverges in adult KO animals as compared to younger KO but not in WT. (A) and (B) Representative raw traces of spiking, superimposed for four trials only for WT younger (black) and KO younger (grey). Scale bar 20mV, 200msec. (C) and (D) Representative Raster plots showing spiking for the step depolarization protocol for WT younger (black) and KO younger (grey). Spikes align precisely and have low within cell variability (CV_w_) across trials for KO younger as compared to WT younger. (E) Bar plot for comparing within cell variability (CV_w_) across multiple trials between WT younger (n=26 cells, black), KO younger (n=19 cells, grey), WT older (n=30 cells, light black) and KO older (n=29 cells, light grey). Spiking variability significantly increases for KO animals with age but not for WT animals (p<0.001, t-test with Bonferroni correction). (F) Bar plot for CV_s500_ for ISI of the step input protocol, a measure of spike precision for all groups as mentioned in (E). Spiking precision significantly decreases for KO animals with age but not for WT animals (p= 0.0002; t-test with Bonferroni correction). (G) Comparison of number of spikes produced during the 900msec stimulus period for each of the groups in (E). There is a non significant change in spike number with age in WT or KO (p=0.6; two way ANOVA) but there was a significant reduction in spike number for KO animals as compared to WT (p=0.021; two-way ANOVA). (H) Superimposed histograms for number of spikes produced for WT older (light grey) and for KO older (dark grey). Solid lines represent Gaussian fitting. The shift in peaks of the histograms shows that there is a greater spread in spike numbers produced in given trials for KO as compared to WT (p<0.001; three way ANOVA). (I) Illustrative f-I curves for a representative cell from older (>6 week) WT animals (black stars), and a cell from older KO animals (filled grey circles). The gain (i.e., slope) for each cell was computed using a regression fit in the roughly linear domain between 50pA to 300pA. (J) Gain computed by fitting a regression line through the f-I curve for each cell as illustrated in (I), for each of the four experimental conditions. The cells were the same as in G. The gains are not significantly different between WT and KO in any age group (n=27 cells for WT older and n=25 cells for KO older; n=26 cells for WT younger and n= 19 cells for KO younger; F _(1, 94)_ = 0.03; p=0.86 for WT and KO differences; two way ANOVA). (K) f-I curve for younger WT (black stars, n=26 cells) vs younger KO (grey plus, 19 cells) CA1 cells. There is no significant difference between WT and KO cells at this age (F _(1, 923)_ = 0.92; p=0.33 for WT and KO differences; two way ANOVA). (L) f-I curve for older WT (black circles, n=27 cells) vs younger KO (grey diamonds, 25 cells) CA1 cells. There is a significant difference between WT and KO cells at this age (F _(1, 1070)_ = 31.03; p<0.001 for WT and KO differences; two way ANOVA).

**Table 1.** Summary of statistical tests.

| Figure Number | Data structure | Type of test | Statistics |
| --- | --- | --- | --- |
| Fig 1E | Non normal | Wilcoxon rank sum test | $p=0.0048$ for $\text{CV}_w$ differences between WT older and KO older. |
| Fig 1F | Non normal | Wilcoxon rank sum test | $p<0.001$ for $\text{CV}_b$ differences between WT older and KO older. |
| Fig 1J | Non normal | 2-way ANOVA | $F_{(1,113)}=5.78$; $p=0.018$ for WT vs KO differences |
| Fig 2E | Normal | 3-way ANOVA | $p<0.001$ for $\text{CV}_w$ differences between WT older and KO older. |
| Fig 2F | Normal | 3-way ANOVA | $p<0.001$ for $\text{CV}_b$ differences between WT older and KO older. |
| Fig 3E | Normal | 2-way ANOVA | $F_{(1,100)} = 4.97$; $p=0.028$ for WT and KO differences; $F_{(1,100)} = 8.05$; $p=0.005$ for interaction factor between age and genotype; $p=0.7$, for WT older vs WT younger; $p<0.001$ for KO older vs KO younger, ttest with bonferroni correction. |
| Fig 3F | Normal | 2-way ANOVA | $F_{(1,99)} = 7.47$, $p=0.007$ for age factor; $F_{(1,99)} = 9.46$; $p=0.003$ for interaction factor between age and genotype; $p=0.0002$ for KO older vs KO younger; $p=0.9$ for WT older vs WT younger; $p=0.0025$ for WT older and KO older |
| Fig 3G | Normal | 2-way ANOVA | $F_{(1,78)} = 5.1$; $p=0.027$ for spike number between WT and KO; $p=0.029$ between WT old and KO old; $p=0.3$ between WT young and KO young. |
| Fig 3J | Normal | 2-way ANOVA | $F_{(1,94)} = 0.03$; $p=0.86$ for WT and KO differences. |
| Fig 3K | Normal | 2-way ANOVA | $F_{(1,923)} = 0.92$; $p=0.33$; WT and KO differences at young age group. |
| Fig 3L | Normal | 2-way ANOVA | $F_{(1,1070)} = 31.03$ $p<0.001$; WT and KO differences at old age group |
| Fig 4B | Normal | 2-way ANOVA | $F_{(1,72)} = 5.35$; $p=0.023$ for difference between WT and KO CA3 cells younger animals |
| Fig 4D | Normal | 2-way ANOVA | $F_{(1,88)} = 1.06$; $p=0.31$ for difference between WT and KO CA1 cells younger animals |
| Fig 4F | Normal | 2-way ANOVA | $F_{(1,75)} = 1.16$; $p=0.28$ for difference between WT and KO CA3 cells older animals. |
| Fig 4H | Normal | 2-way ANOVA | $F_{(1,59)} = 7.42$; $p=0.009$ for difference between WT and KO CA1 cells for older animals. |
| Fig 5C | Normal | 2-way ANOVA | $F_{(1, 46)} = 4.8$; $p=0.033$ for mAHP WT and KO differences. $p=0.2$ between WT younger and KO younger; $p=0.017$ for WT older and KO older. |
| Fig 5D | Normal | 2-way ANOVA | $F_{(1, 54)} = 8.84$; $p=0.004$ for sAHP WT and KO differences. $p=0.07$ between WT younger and KO younger; $p=0.008$ for WT older and KO older. |
| Fig 5F | Non normal | Wilcoxon rank sum test | $p=0.91$ for $I_h$ % sag between WT and KO |
| Fig 5H | Normal | 2-way ANOVA | $F_{(1, 258)} = 2.54$; $p=0.11$ between WT and KO for $I_M$ current |
| Fig 6C | Normal | 2-way ANOVA | $F_{(1, 240)} = 33.3$; $p<0.001$ for WT older and KO older SK currents |
| Fig 6D | Normal | 2-way ANOVA | $F_{(1, 189)} = 0.51$; $p=0.5$ for WT younger and KO younger SK currents |
| Fig 6E | Normal | 2-way ANOVA | $F_{(1, 229)} = 0.01$; $p=0.9$ for WT older and WT younger SK currents |
| Fig 6F | Normal | 2-way ANOVA | $F_{(1, 209)} = 21.02$; $p<0.001$ for KO older and KO younger SK currents |
| Fig 7B | Normal | 2-way ANOVA | $F_{(1, 24)} = 0.48$; $p=0.5$ for WT younger vs KO younger for CA3 |
| Fig 7C | Normal | 2-way ANOVA | $F_{(1, 24)} = 0.21$; $p=0.6$ for WT younger vs KO younger for CA1 |
| Fig 7D | Normal | 2-way ANOVA | $F_{(1, 24)} = 0.3$; $p=0.6$ for WT older vs KO older for CA3 |
| Fig 7E | Normal | 2-way ANOVA | $F_{(1, 24)} = 0.25$; $p=0.61$ for WT older vs KO older for CA1 |
| Fig 8E | Normal | 2-way ANOVA | $F_{(1, 48)} = 0.96$ for effect of apamin; $F_{(1, 48)} = 0.004$ for WT and KO differences |
| Fig 8F | Normal | 2-way ANOVA | $F_{(1, 96)} = 157.5$; $p<0.001$ for WT and KO differences; $F_{(1, 96)} = 53.1$; $p<0.001$ for effect of apamin on $CV_b$ |
| Fig 8G | Normal | 2-way ANOVA | $F_{(1, 49)} = 9.63$; $p=0.003$ for WT and KO differences; $F_{(1, 49)} = 2.87$; $p=0.09$ for effect of apamin; $p=0.006$ for WT pre apamin and KO pre apamin; $p=0.05$ for KO pre apamin and KO post apamin; $p=0.23$ WT pre apamin and KO post apamin. |
| Fig 8H | Normal | 2-way ANOVA | $F_{(1, 81)} = 2.7$; $p=0.1$ for differences between WT and KO; $F_{(1, 81)} = .40.5$ $p<0.001$ for affect of BAPTA |

We performed a 3-way ANOVA in which the factors were: across trials, across cells, and WT vs. KO for older age group data. The ANOVA results were highly significant for variability in spike numbers between cells (CV_b_) (F _(29, 1420)_ = 32.04, p<0.001) and differences in spike numbers between WT and KO for older animals (F _(1, 1420)_ = 209.9, p<0.001) but across-trial comparisons did not exhibit a significant difference (F _(24, 1420)_ = 0.15, p=1).Thus, there is increase in spike variability with lowered spiking precision for adult KO mice as compared to juvenile KO mice. This divergence which is observed for KO animals with age does not happen for WT animals.

### Differential effect of FMRP KO in different brain regions

Mouse models of FXS are known to exhibit increased excitability (Contractor et al., 2015; Deng and Klyachko, 2016; Luque et al., 2017). In contrast to previous studies, we observed that excitability was reduced in CA1 neurons in adult mice but not in juveniles. Deng et. al. 2019 have observed increased CA3 excitability in juveniles which was in contrast to what is we observed in CA1, namely, that there is no significant effect on excitability of CA1 pyramidal cells at this time point. In order to address this contrasting result, we repeated the protocol from Deng et. al. in our FXS model, and compared outcomes between CA3 and CA1 (Figure 4). In agreement with their study we observed significantly increased spiking in CA3 cells of FXS mice for some of the input holding currents (# of spikes in 60 s for −48mV holding potential for WT 131.9 ± 17.42, n=10 cells; # of spikes in 60 s for −48mV holding potential for KO 186.3 ± 27.27, n=10 cells; F _(1, 72)_ = 5.35, p=0.023 for difference between WT and KO; two-way ANOVA) (Figure 4A, 4B). We did similar experiments in CA1 pyramidal cells, using the same protocol as in Figure 4A and B. We did not find any difference in excitability between WT and KO (# of spikes in 60 s for −48mV holding potential for WT 355.58 ± 40.93, n=12 cells; # of spikes in 60 s for −48mV holding potential for KO 358.67 ± 22.56, n=12 cells; F _(1, 88)_ = 1.06, p=0.31 for difference between WT and KO, two-way ANOVA) (Figure 4C, 4D). Upon seeing this differential effect of FMRP KO in different regions of hippocampus, we were interested to see if this might be a function of age. Therefore, we recorded similar protocols from CA3 and CA1 of older animals. We found that the hyper-excitability seen in younger CA3 cells, goes away, leading to no significant difference in spiking activity in these cells for WT vs KO (# of spikes in 60 s for −48mV holding potential for WT 313.6 ± 83.8, n=10 cells; # of spikes in 60 s for −48mV holding potential for KO 181.1 ± 54.9, n=10 cells; F _(1, 75)_ = 1.16, p=0.28 for difference between WT and KO, two-way ANOVA) (Figure 4E, 4F). In agreement with our previous results as shown in Fig 3, we observed reduced excitability in CA1 KO cells as compared to WT (# of spikes in 60 s for −50mV holding potential for WT 135.6 ± 35.2, n=8 cells; # of spikes in 60 s for −50mV holding potential for KO 54 ± 20.4, n=8 cells; F _(1, 59)_ = 7.42, p=0.009 for difference between WT and KO, two-way ANOVA) (Figure 4G, 4H). Using these experiments, we were able to replicate previous published results of Deng et. al. 2019. In addition, we found that there were contrasting and age-dependent effects of FMRP KO on the spiking properties of CA1 and CA3 pyramidal neurons.

**Figure 4.**
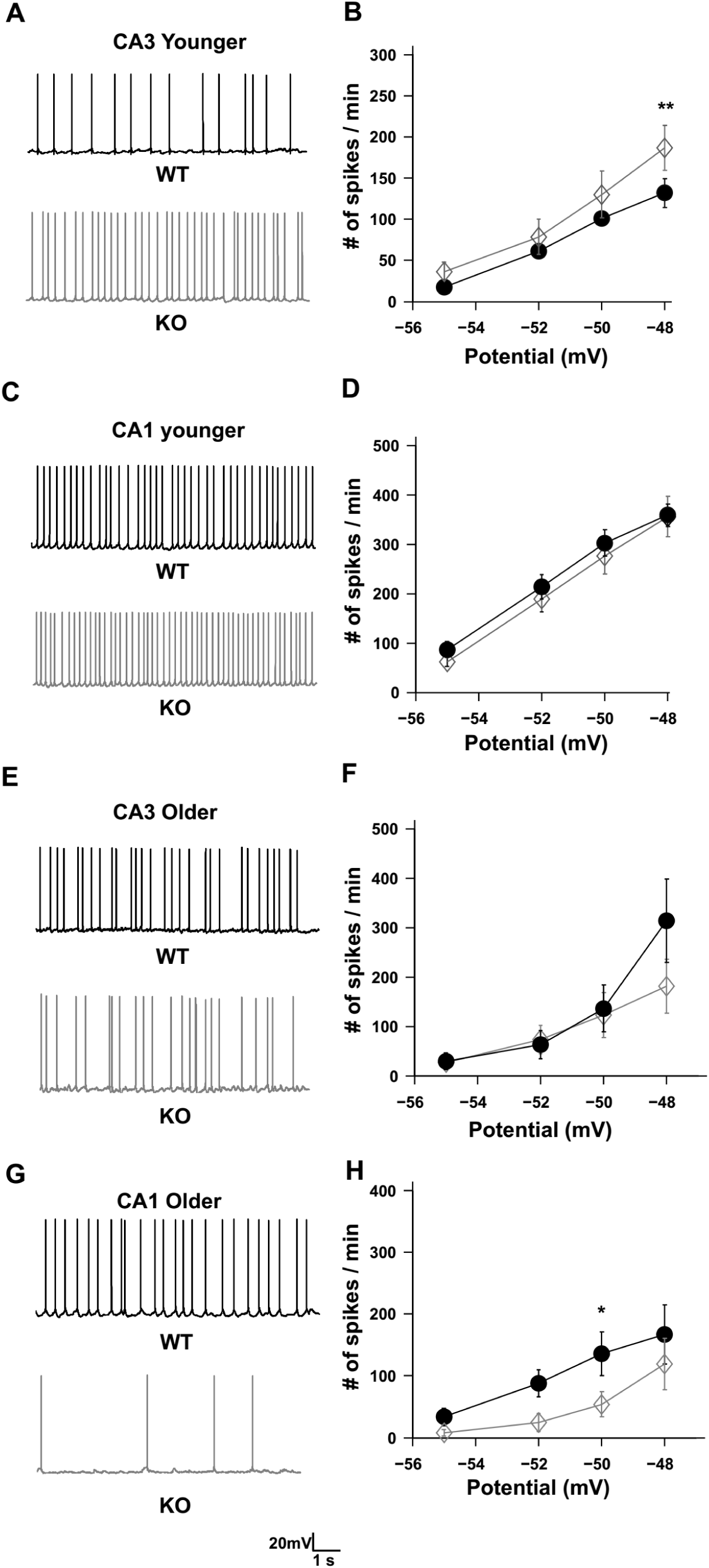
Contrasting effects of FMRP KO in different regions of hippocampus. A) Representative traces showing increased excitability between WT (black) and KO (grey) CA3 cells in younger animals. Traces are from the recording where the cells are held at −48mV. Scale bar 20mV, 1s. B) Line plot showing increased excitability for KO CA3 cells (n=10 cells, grey diamond) as compared to WT CA3 cells (n=10 cells, filled black circles), when the cells were held at −48mV, for younger animals. For other holding potentials the spike numbers are not significantly increased for KO cells (F _(1, 72)_ = 5.35, p=0.023 for difference between WT and KO, two way ANOVA, error bar represent SEM) C) Representative traces showing no change in excitability between WT (black) and KO (grey) CA1 cells in younger animals. Traces are from the recording where the cells are held at −48mV. Scale bar 20mV, 1s. D) Line plot showing no change in excitability for KO CA1 cells (n=12 cells, grey diamond) as compared to WT CA1 cells in younger animals (n=12 cells, filled black circles) (F _(1, 88)_ = 1.06, p=0.31 for difference between WT and KO, two-way ANOVA, error bar represent SEM). E) Representative traces showing no change in excitability between WT (black) and KO (grey) CA3 cells in older animals. Traces are from the recording where the cells are held at −48mV. Scale bar 20mV, 1s. F) Line plot showing no change in excitability for KO CA3 cells (n=10 cells, grey diamond) as compared to WT CA3 cells (n=10 cells, filled black circles), when the cells were held at −48mV, for older animals. (F _(1, 72)_ = 1.17, p=0.28 for difference between WT and KO, two way ANOVA, error bar represent SEM) G) Representative traces showing reduced excitability for KO cells (grey) as compared to (black) and KO CA1 cells in older animals. Traces are from the recording where the cells are held at −50mV. Scale bar 20mV, 1s. H) Line plot showing decreased excitability for KO CA1 cells (n=8 cells, grey diamond) as compared to WT CA1 cells (n=8 cells, filled black circles), when the cells were held at −60mV, for older animals. For other holding potentials the spike numbers are not significantly decreased for KO cells (F _(1, 59)_ = 7.42, p=0.009 for difference between WT and KO, two way ANOVA, error bar represent SEM)

### mAHP currents are elevated in older KO animals as compared to WT but no significant changes in I_h_ and M currents

We next investigated the mechanisms for increased variability in KO animals, on the basis of the above observations of increased variability and fewer spikes (Fig 3G and 3L) in the later part of the current pulse. In particular, we hypothesized that medium afterhyperpolarization (mAHP) and slow afterhyperpolarization (sAHP) currents were likely candidates to mediate such slower-onset effects. We asked if these currents might be altered in FXS cells.

To test this hypothesis we compared AHPs for WT vs. KO in older and younger animals. Cells were held at −70mV to record AHPs. We quantified AHPs by examining the hyperpolarization elicited with a spike train of 5 spikes in 100msec. Hyperpolarization was quantified at mAHP peak (50msec after the spike train) and at sAHP peak (200msec after the spike train). We found that mAHP was elevated only in older KOs but not in younger KOs when compared to their WT littermates (mAHP for WT older −0.26 ± 0.16, n=12 cells; mAHP for KO older −0.71 ± 0.15, n=13 cells; mAHP for WT younger 0.41 ± 0.2, n=14 cells; mAHP for KO younger −1.14 ± 0.5, n=18 cells; F_(1, 46)_ = 4.8, p=0.033 for WT and KO differences) (Fig 5C, detailed statistics shown in table 1). We found similar results for sAHP measurements as well. We found that sAHP was significantly elevated only for older KO animals as compared to their WT littermates but not significantly different for younger KO animals as compared to their younger WT littermates (sAHP for WT older 0.006 ± 0.1, n=12 cells; sAHP for KO older −0.37 ± 0.11, n=13 cells; sAHP for WT younger 0.04 ± 0.12, n=14 cells; sAHP for KO younger 0.047 ± 0.2, n=18 cells; F _(1, 54)_ = 8.84, p=0.004 for WT and KO differences) (Fig 5D). As there was no significant difference in spike variability and spike time precision in younger KO animals, we hypothesized that the observed spiking variability in older KO animals might be due to elevated mAHPs, which has a strong change with age in KO animals. Further, the stimulus period used in the previous experiments (Fig 1) was of 900ms which is the peak time of activation for mAHP mediating ion channels.

**Figure 5.**
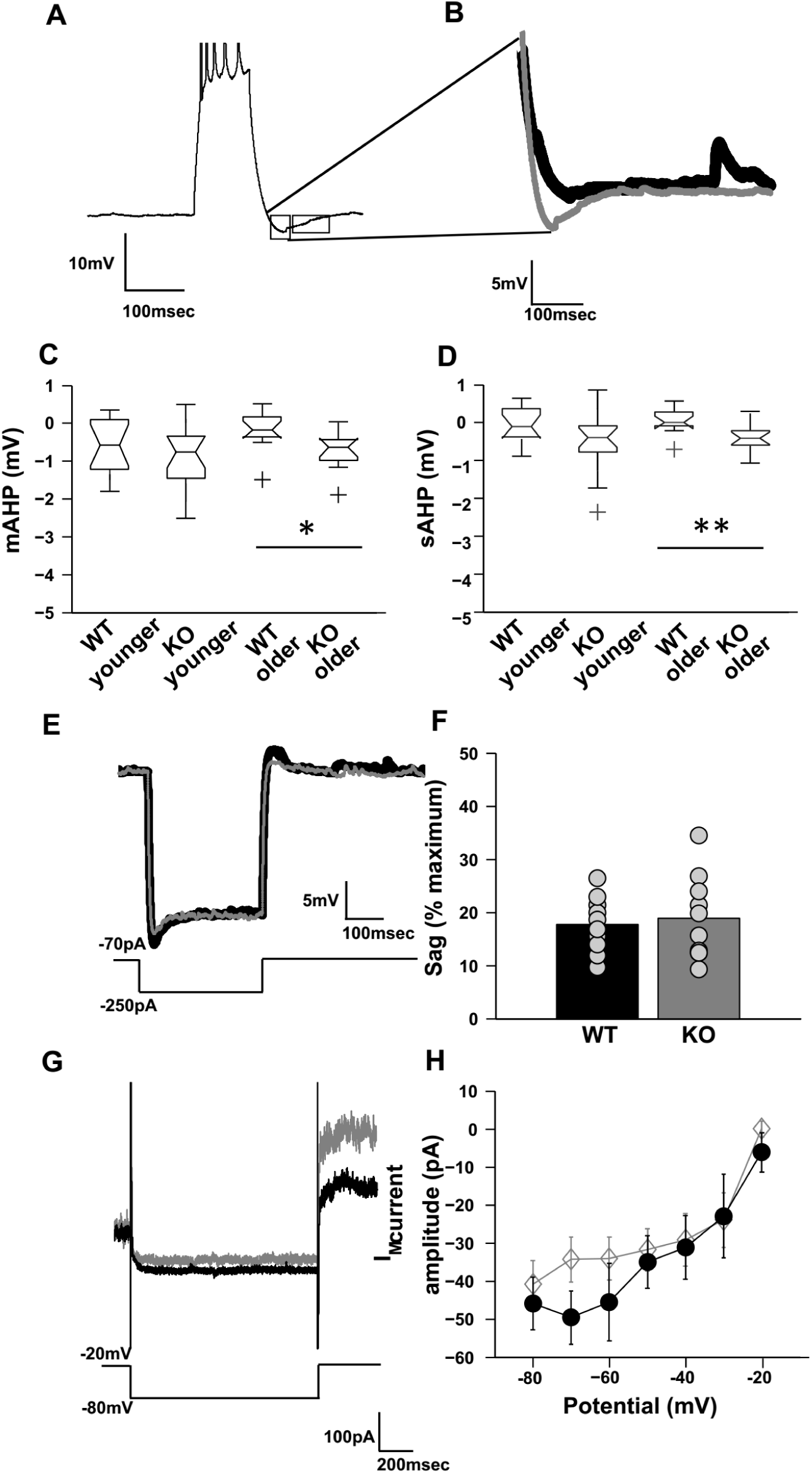
mAHP and sAHP are elevated in FXS KO neurons. (A) Representative trace for the protocol to measure mAHP and sAHP. Hyperpolarization within 50msec of the spike train (1) is quantified as mAHP while hyperpolarization following the spike train 200msec after the spike train (2) is quantified as sAHP. Scale bar 10mV, 100msec. (B). Superimposed traces of WT older and KO older AHP currents showing elevated mAHP currents for KO cell (grey) as compared to WT cell (black). Scale bar 5mV, 100msec. (C) Box plot showing mAHP currents for all groups (n=12 cells for WT older, n=13 cells for KO older, n=14 cells for WT younger and n=18 cells for KO younger). mAHP are elevated only for older KO animals vs. older WT (p=0.018; t-test with Bonferroni correction, error bar represent SEM). (D) Box plot showing sAHP currents for all groups (n=12 cells for WT older, n=13 cells for KO older, n=14 cells for WT younger and n=18 cells for KO younger). sAHP is elevated for older KO vs older KO (p=0.008; t-test with Bonferroni correction) but not in younger KO vs younger WT animals as well (p=0.06 for sAHP between WT younger and KO younger, error bar represent SEM). (E) Representative trace for the protocol to measure I_h_ current. Superimposed traces of WT (black) and KO (grey) showing no significant difference in the response. Sag produced in response to 250pA hyperpolarizing pulse was quantified to estimate I_h_ current. Scale bar 5mV, 100msec. (F) Bar plot for showing % sag produced due to I_h_ current for WT older and KO older (n=10 cells for both WT and KO older). There was no significant difference in sag due to I_h_ current (p=0.91, Wilcoxon’s test). (G) Representative traces showing XE991 sensitive M current for WT older (black) and KO older (grey). The traces were plotted by finding the difference between pre and post blocker current trace for input step from −20mV to −80mV. Scale bar 100pA, 200msec. (H) Line plot showing IV curve of XE 991 sensitive M current. There is no significant difference in the M currents between WT older (filled black circles) and KO older (grey diamond) for any of the input steps. (n=19 cells for both WT and KO, F _(1, 258)_ = 2.54, p=0.11 between WT and KO, two-way ANOVA, error bars represent SEM)

The mAHP in CA1 cells is mediated by I_h_, M and SK currents. A previous study has shown that I_h_ currents are not significantly altered in soma of FXS cells (Brager et al., 2012). M currents are known to affect the Resting Membrane Potential (RMP) and spike adaptation index (Madison et al., 1987; Guan et al., 2011; Nigro et al., 2014; Hönigsperger et al., 2015). Neither RMP nor spike adaptation index were significantly changed in FXS cells, indicating that M currents might not be responsible for the observed phenotype. To further test this idea, I_h_ currents and M currents were recorded from soma of CA1 cells and compared between WT and KO for adult animals. As there was no difference in mAHP at juvenile stage for KO cells, we did not measure I_h_ and M currents for them. I**_h_** channel activity was recorded using a hyperpolarization pulse. The voltage sag produced in response to the pulse, was quantified according to a method adapted from Brager and Johnston, 2012 (Methods, Fig 5E). As observed previously, there was no significant difference in I**_h_** sag between WT and KO CA1 cells, at somatic level. (% sag for WT 17.7 ± 1.64, n=10 cells; % sag for KO 18.87 ± 2.51, n=10 cells; p=0.91; Wilcoxon rank sum test) (Fig 5F).

To measure M currents we used the protocol adapted from Shah et al 2002 (Methods). According to the protocol the cells were held at −20mV and M currents were deactivated in steps from −20mV to −80mV. M current specific blocker XE991 was used to obtain the post blocker trace. Post blocker trace was subtracted from pre blocker trace to obtain XE991 sensitive M currents (Fig 5G). We found no significant differences in M currents between WT and KO CA1 cells. (Mean M current for step −80mV for WT −40.9 ± 6.33, n=19 cells; Mean M current for step - 80mV for KO 45.94 ± 6.87, n=19 cells; F _(1, 258)_ = 2.54, p=0.11 between WT and KO, two-way ANOVA) (Fig 5H). Overall, these observations suggest that neither I_h_ nor M currents contribute to the observed elevation in mAHP in KO animals.

### SK currents are elevated in older KO animals as compared to WT

SK currents are also known to contribute substantially to mAHP (Sah and Clements, 1999, Stocker et. al 1999). It has also been shown that Ca2+ influx from VGCCs is elevated in FXS cells (Deng et al., 2011, 2013), indicating elevated Ca2+ levels intracellularly and hence increased SK channel activation. Thus, we hypothesized that SK currents are elevated in FXS cells, leading to the observed elevated mAHP in these cells. To test this, we used a procedure from Carlen, 1997. Outward currents were evoked using step depolarization of the cell from −55mV to +5mV and above (Fig 6A and 6B), in voltage clamp mode. Apamin was bath applied to obtain post apamin trace. The difference current of pre apamin and post apamin was quantified as SK currents. These currents were significantly higher in KO animals as compared to WT for adult age group (SK current for older WT at +15mV step 54.2 ± 9, n=13 cells; SK current for older KO at +15 mV step 131.73 ± 25.5, n=13 cells; F_(1, 240)_ = 33.3, p<0.001 between WT and KO, two-way ANOVA)(Fig 6C). However, this phenotype was only seen in adult WT vs adult KO animals. When similar experiments were done in juveniles of both genotype, there was no significant difference in SK currents (SK current for younger WT at +15mV step 61.93 ± 11.9, n= 11 cells; SK current for younger KO at +15mV step 51.2 ± 8.9, n=9 cells; F_(1, 189)_ = 0.51, p=0.5, two way ANOVA) (Fig 6D). To delineate the developmental change in SK channels levels between WT and KO, comparisons were made within the same genotype, between different age groups. There was no significant change in the levels of SK currents with age for WT animals (SK current for Older WT at +15mV step 54.2 ± 9, n=13 cells; SK current for younger WT at +15mV step 61.93 ± 11.9, n= 11 cells; F_(1, 229)_ = 0.01, p=0.9 two way ANOVA) (Fig 6E). On the contrary there was a significant increment in levels of SK currents for KO animals as they progressed from juvenile to adult stage (SK current for older KO at +15 mV step 131.73 ± 25.5, n=13 cells; SK current for younger KO at +15mV step 51.2 ± 8.9, n=9 cells; F_(1,209)_=21.02, p<0.001; two-way ANOVA) (Fig 6F). Together, these observations show that there is an increase in levels of SK currents with age in KO animals, and we note that this correlates with the worsening of spike reliability between adult and juvenile KO animals as described in Fig 3.

**Figure 6.**
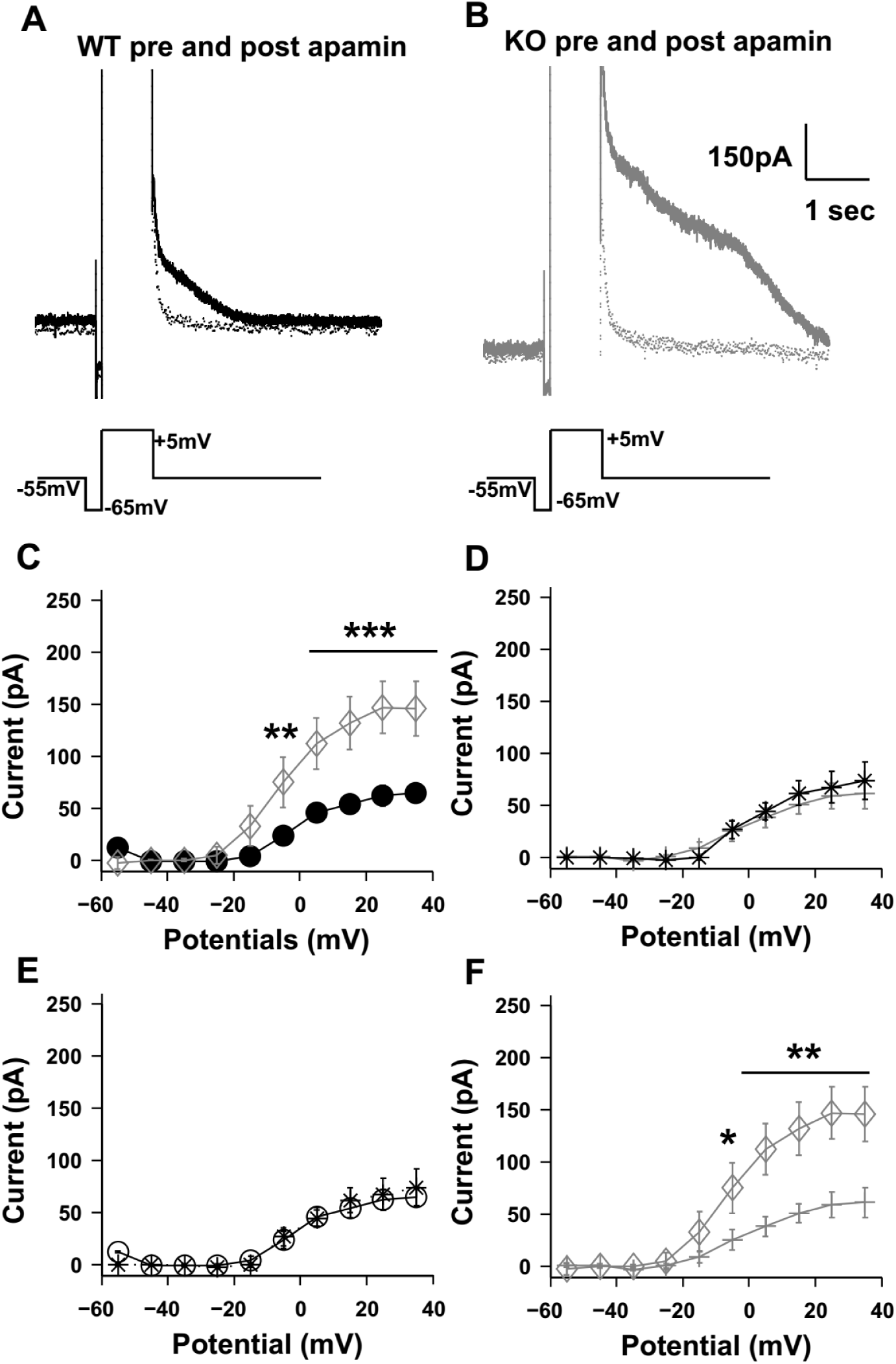
Apamin sensitive SK currents are elevated in KO neurons as compared to WT. (A) Representative traces showing the outward currents elicited by step depolarization voltage commands in Voltage Clamp, for WT neuron (black) and (B) for KO neuron (grey), pre apamin (solid line) and post apamin (dotted line) perfusion. Trace below indicates command voltage protocol. Scale bar 150pA, 1sec. (C) IV curve of the apamin sensitive current elicited across multiple steps for WT older cells (n=13 cells, filled black circles) and KO older cells (n=13 cells, grey diamond). Currents are significantly elevated for KO cells as compared to WT cells, for multiple steps. (F _(1, 240)_ = 33.3, p<0.001 between WT and KO, two-way ANOVA, error bars represent SEM) (D) IV curve of the apamin sensitive current elicited across multiple steps WT younger cells (n=11 cells, black star) and KO younger cells (n=9 cells, grey plus). Currents between KO and WT cells at this age group were not significantly different. (F _(1, 189)_ = 0.51; p=0.5; two way ANOVA, error bars represent SEM). (E) IV curve of the apamin sensitive current elicited across multiple steps for WT younger cells (n=11 cells, black stars) and WT older cells (n=13 cells, black circles). There is no age dependent change in SK currents for WT animals (F _(1, 229)_ = 0.01, p=0.9 two way ANOVA, error bars represent SEM). (F) IV curve of the apamin sensitive current elicited across multiple steps for KO younger cells (n=9 cells, grey plus) and KO older cells (n=13 cells, grey diamonds). There is an age dependent increase in SK currents in KO animals (F _(1, 209)_ = 21.02, p<0.001; two-way ANOVA, bars represent SEM).

### SK channel expression is unchanged in CA1 and CA3 regions for KO as compared to WT

We have observed that SK current levels are elevated in KO CA1 cells as compared to WT (Fig 6). This elevation might arise due to increase in the levels of SK channels or due to change in SK channel kinetics. The latter might arise due to change in the kinetics of the SK channel itself or due to changes in activity of other channels which have an effect on SK channels. The putative candidates might be Ca^2+^ channels known to be effected in FXS (Meredith et al., 2007; Ferron et al., 2014,Brown et al., 2001; Contractor et al., 2015). Previous published results have not shown SK channels to be targets of FMRP for transcription regulation, decreasing the likelihood that the levels will be affected in the CA1 region (Darnell et al., 2011; Deng et al., 2018). To specifically test this point, immunofluorescence experiments were used to assess the expression of SK channels. We performed in-situ immunolabeling to measure the intensity of the SK2 isoform in CA3 and CA1 regions of the hippocampus using a specific antibody (Methods) (Fig 7A). We analyzed levels of SK2 protein in different regions and sub sections of the hippocampus for both WT and KO. Similar experiments were done on both adult and juvenile slices. In juvenile animals, we have seen increased number of spikes in CA3 with no significant change in CA1 (Fig 4). However, the levels of SK2 protein were not significantly altered in KO slices for either CA3 (N=4 animals for WT; N=4 animals for KO; F _(1, 24)_ = 0.48; p=0.5 for WT vs KO for CA3; two way ANOVA) (Fig 7B) or CA1 (N=4 animals for WT; N=4 animals for KO; F _(1, 24)_ = 0.21; p=0.6 for WT vs KO for CA1; two way ANOVA) (7C). For adult animals we have seen reduction in spike number in CA1 region (Fig 3G and 3H), but no changes in SK2 levels could be found in either CA1 or CA3 (CA1: N=4 animals for WT; N=4 animals for KO; F _(1, 24)_ = 0.25; p = 0.61 for WT vs KO for CA1; two way ANOVA) (Fig 7E) (CA3: N=4 animals for WT; N=4 animals for KO; F _(1, 24)_ = 0.3; p=0.6 for WT vs KO for CA3; two way ANOVA) (Fig 7D). Overall, these labeling studies suggest that the KO does not affect SK expression in either CA1 or CA3. The latter observation is in agreement with previous results (Deng et al., 2018).

**Figure 7.**
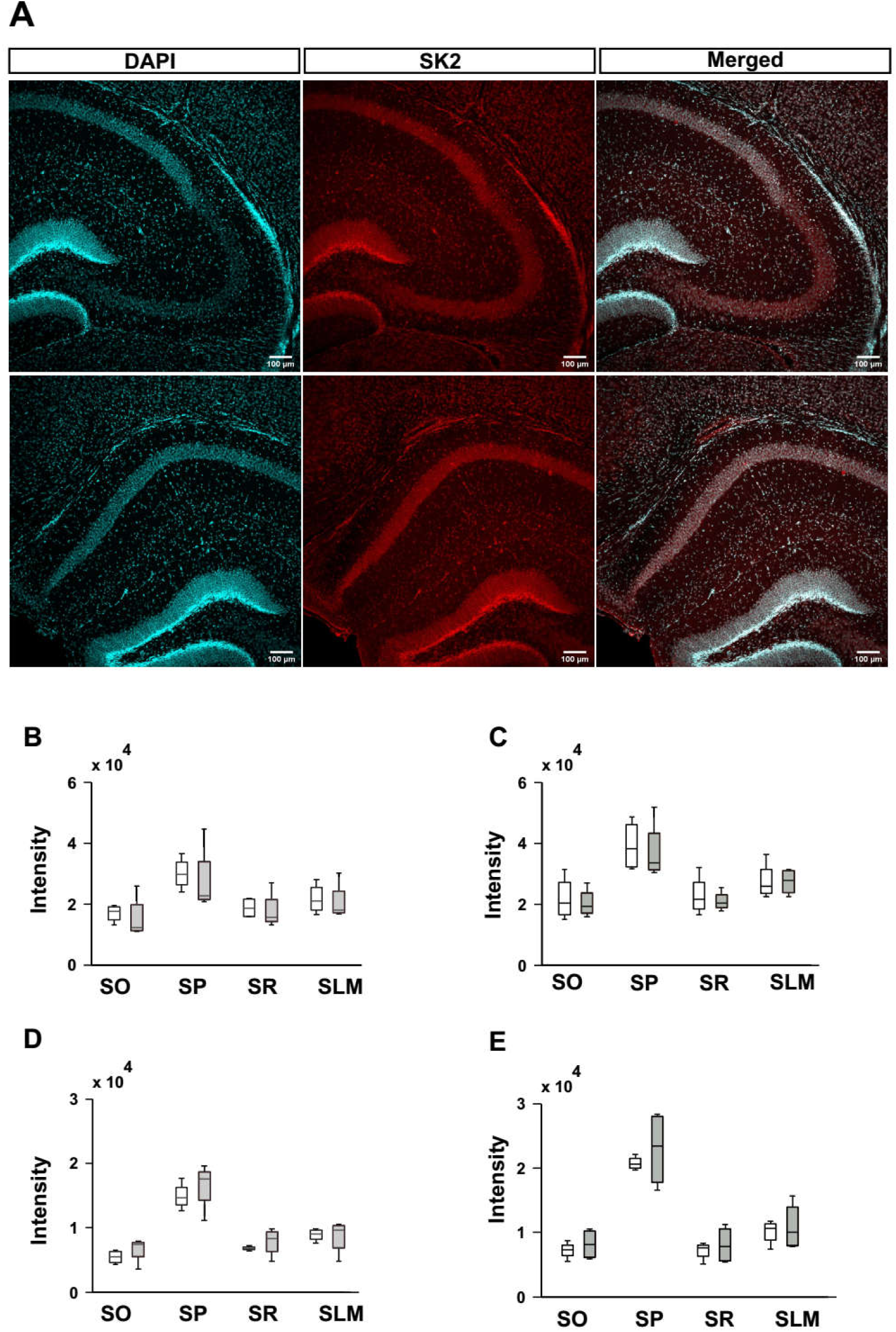
There is no significant change in SK2 channel levels in KO as compared to WT. A. Representative images of SK2 labeling in CA3 (upper panel) and CA1 (lower panel) for a WT animal. Green false color used for DAPI for better visualization. Scale bar 100µm. B. Analyzed box plot for mean intensity of SK2 channel in CA3 for juvenile animals. (N=4 animals for WT; N=4 animals for KO; F _(1, 24)_ = 0.48; p=0.5 between WT and KO; two way ANOVA, error bars represent SEM) C. Analyzed box plot for mean intensity of SK2 channel in CA1 for juvenile animals. (N=4 animals for WT; N=4 animals for KO; F _(1, 24)_ = 0.21; p=0.6 between WT and KO for CA1; two way ANOVA, error bars represent SEM) D. Analyzed box plot for mean intensity of SK2 channel in CA3 for adult animals. (N=4 animals for WT; N=4 animals for KO; F _(1, 24)_ = 0.3; p=0.6 between WT and KO for CA1, error bars represent SEM) E. Analyzed box plot for mean intensity of SK2 channel in CA1 for adult animals. (N=4 animals for WT; N=4 animals for KO; F _(1, 24)_ = 0.25; p=0.61 between WT and KO for CA1; two way ANOVA, error bars represent SEM)

### Apamin partially rescues the phenotype of increased variability in KO animals

Having shown that elevated and variable SK currents play a role in cellular firing variability, we then asked if blockage of the SK current might rescue the phenotype of increased variability. We observed that WT and KO populations responded differently upon blocking SK currents using apamin. In WT cells, the mean of within cell variability (CV_w_) between trials increased (CV_w_ for WT cells pre apamin 0.03 ± 0.002, n=13 cells; CV_w_ for WT cells post apamin 0.04 ± 0.005, n=13 cells) (Fig 8A,8B and 8E). However, for KO cells mean of within cell variability (CV_w_) between trials decreased (CV_w_ for KO cells pre apamin 0.06 ± 0.008, n=13 cells; CV_w_ for KO cells post apamin 0.05 ± 0.009, n=13 cells) (Fig 8C,8D and 8E). Apamin treatment had no significant affect on within-cell variability (CV_w_) for either WT or KO. However, it leads to a partial rescue such that post apamin KO CV_w_ values were not significantly different from pre apamin WT CV_w_ values. A complete rescue by apamin would have resulted in a significant reduction of the CV_w_ of KO cells (which did not happen) as well as a convergence of variability between KO and WT (which did) (F _(1, 48)_ = 0.96 for effect of apamin; F _(1, 48)_ = 0.004 for WT and KO differences; two-way ANOVA) (Fig 8E). We performed similar analysis for between cell (CV_b_) variability. Apamin perfusion led to a partial rescue in CV_b_ parameter as well such that there was a significant reduction in CV_b_ of KO cells upon apamin perfusion, but no convergence of variability between WT and KO (CV_b_ for WT cells pre apamin 0.414 ± 0.002; CV_b_ for WT cells post apamin 0.413 ± 0.002; CV_b_ for KO cells pre apamin 0.47 ± 0.003; CV_b_ for KO cells post apamin 0.43 ± 0.002; F _(1, 96)_ = 157.5; p<0.001 for WT and KO differences; F _(1, 96)_ = 53.1; p<0.001 for effect of apamin on CV_b;_ two-way ANOVA)(Fig 8F). CV_s500_ also showed a partial rescue with apamin perfusion (CV_s500_ for WT cells pre apamin 0.042 ± 0.007; CV_s500_ for WT cells post apamin 0.042 ± 0.005; CV_s500_ for KO pre apamin 0.09 ± 0.03; CV_s500_ for KO post apamin 0.04 ± 0.007; F _(1, 49)_ = 9.63; p=0.003 for WT and KO differences; F _(1, 49)_ = 2.87; p=0.09 for effect of apamin; two-way ANOVA) (Fig 8G). Overall, we find that blockage of SK currents by apamin partially rescues the variability phenotypes, consistent with our interpretation that elevation in SK currents has a role in variability.

**Figure 8.**
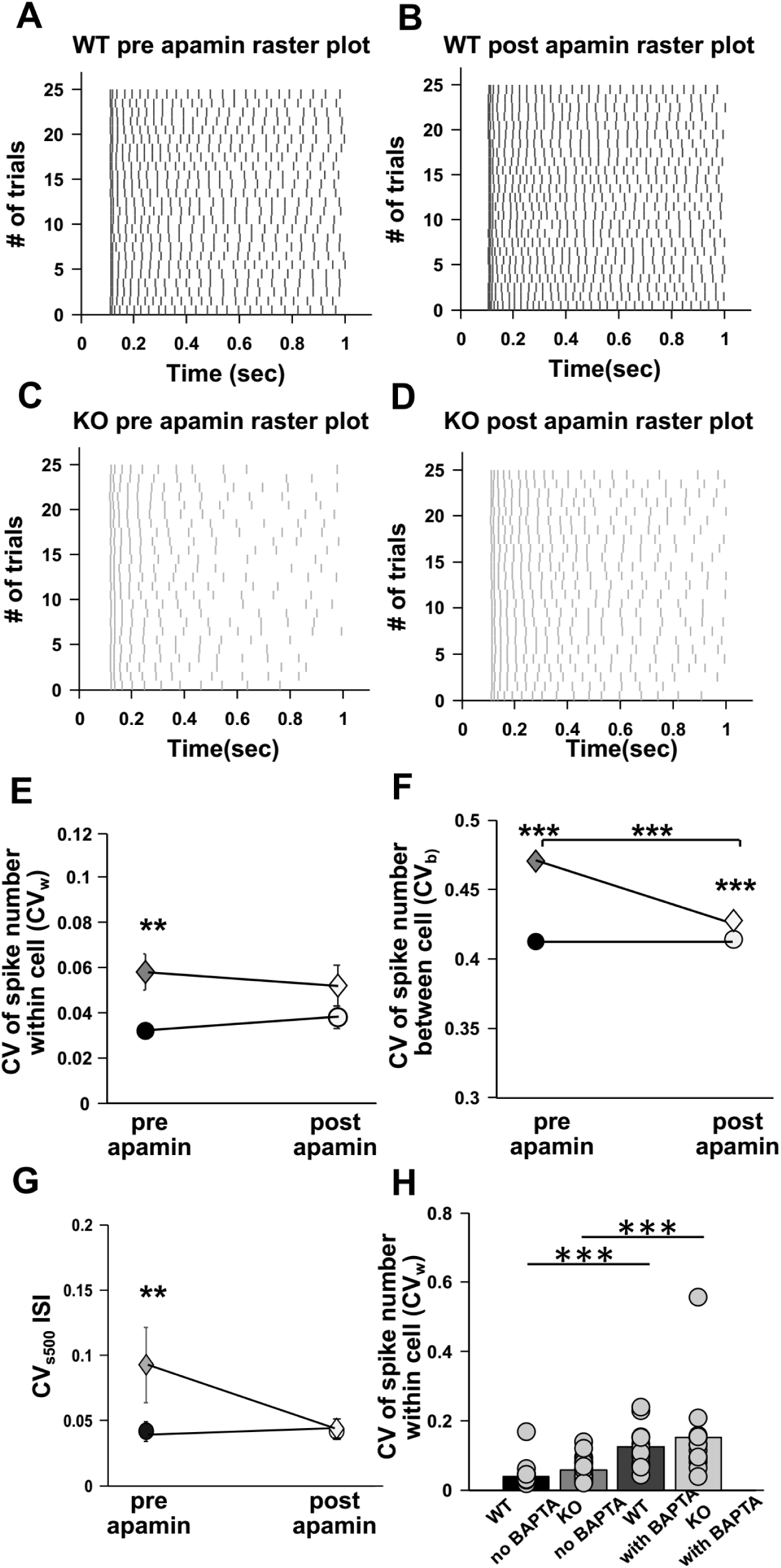
Partial rescue of within cell variability (CV_w_), between cell variability (CV_b_) and spike imprecision in KO neurons upon apamin perfusion. (A) Representative Raster plot showing spiking for step depolarization protocol for WT, pre apamin. (B) Representative Raster plot showing spiking for step depolarization protocol for WT, post apamin. Same cell as A. (C) Representative Raster plot showing spiking for step depolarization protocol for KO, pre apamin. (D) Representative Raster plot showing spiking for step depolarization protocol for KO, post apamin. Same cell as C. (E) Change in within cell variability parameter (CV_w_) for pre (filled black circles) and post apamin perfusion in WT (n=13 cells, empty black circles) and in KO (n=13 cells, filled grey diamonds for pre apamin and empty grey diamonds for post apamin). There is a partial rescue in CV_w_ parameter post apamin for KO cells vs. WT cells (F _(1, 48)_ = 0.96 for effect of apamin; F _(1, 48)_ = 0.004 for WT and KO differences; 2-way ANOVA with replication, error bars represent SEM) (F) Plot showing change in between cell variability parameter (CV_b_) for pre (filled black circles) and post apamin perfusion WT (n=13 cells, empty black circles) and for pre and post apamin perfusion KO (n=13 cells, filled grey diamonds for pre apamin and empty grey diamonds for post apamin). There is a partial rescue in CV_b_ parameter for KO cells vs. WT cells (F _(1, 96)_ = 157.5; p<0.001 for WT and KO differences; F _(1, 96)_ = 53.1; p<0.001 for effect of apamin on CV_b;_ two-way ANOVA with replication error bars represent SEM). (G) Plot for CV_s500_ for ISI, a measure of spike precision for pre (filled black circles) and post apamin perfusion WT (n=13 cells, empty black circles) and for pre and post apamin perfusion KO (n=13 cells, filled grey diamonds for pre apamin and empty grey diamonds for post apamin). There is a partial rescue of CV_s500_ parameter upon apamin perfusion (F _(1, 49)_ = 9.63; p=0.003 for WT and KO differences; F _(1, 49)_ = 2.87; p=0.09 for effect of apamin; two-way ANOVA error bars represent SEM). (H) Bar plot showing effect of BAPTA on within cell variability (CV_w_) for WT (n=30 cells without BAPTA and n=12 cells with BAPTA) and KO (n=29 cells without BAPTA and n=12 cells with BAPTA). Chelating Ca2+ using BAPTA, lead to a significant increase in spike variability within cell (CV_w_) for both WT and KO cells (F _(1, 81)_ = 2.7; p=0.1 for differences between WT and KO; F _(1, 81)_ = 40.5 p<0.001 for affect of BAPTA; 2-way ANOVA with replication, error bars represent SEM).

**Figure 9.**
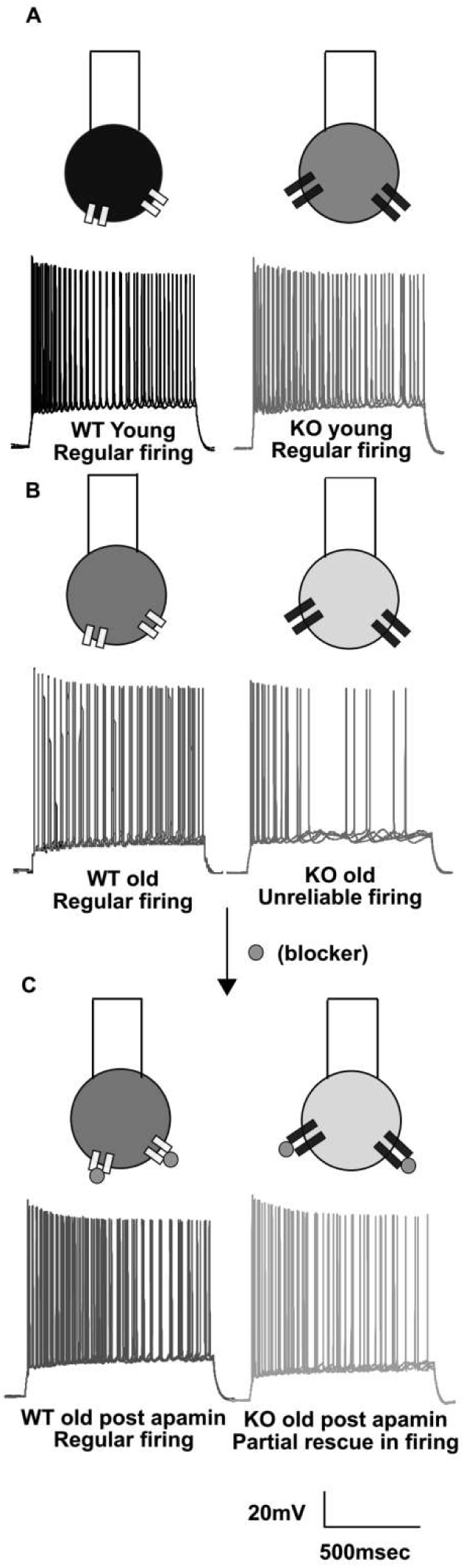
Schematic summary plot indicating developmental increase in spiking variability and the role of SK channels. All spiking traces are actual raw data and show superimposed traces for 4 trials in WT (Black) and KO (Grey) respectively. (A) Schematic of neurons, showing Younger WT (Black) and Younger KO (Grey). In young animals we observe precise and reliable spiking across multiple trials. Scale bar 20mV, 500msec. (B) Schematic of Older WT and Older KO neurons. KO neurons have larger SK currents which are represented by larger icons for SK channels. Noticeably, KO spikes are more variable and less precise especially in the later part of the trial. (C) Apamin (shown as small grey circles) is used to block SK channels present on soma. Upon apamin perfusion spiking in WT (black) becomes more variable whereas spiking in KO (light grey), increases in frequency leading to increased spike precision and reliability.

### Blocking Calcium has no rescue affect

From the above observations, blocking SK currents led to only a partial rescue in the phenotype of increased imprecision and unreliability for spikes in KO cells. We therefore asked if Ca^2+^, as an upstream modulator of SK, could also contribute to a partial rescue. To do this we used BAPTA to chelate intracellular Ca^2+^. We added BAPTA (10mM) into the intracellular recording pipette and assessed spike precision 10 minutes after patching. We found that chelating calcium using BAPTA led to further increase in spike variability instead of rescue. Spike reliability within cell was measured (CV_w_) in these experiments and it was found that spikes became significantly more variable post BAPTA than pre BAPTA for both WT and FXS cells (CV_w_ for WT cells without BAPTA 0.038 ± 0.005, n=30 cells; CV_w_ for WT cells with BAPTA 0.12 ± 0.02, n=12 cells; p<0.001; CV_w_ for KO cells without BAPTA 0.057 ± 0.006, n=29 cells; CV_w_ for KO cells with BAPTA 0.16 ± 0.04; F _(1, 81)_ = 40.5 p<0.001 for affect of BAPTA; two way ANOVA) (Fig 8H). Since Ca^2+^ has a large number of ion-channels as other targets, we concluded that it was not a good target for rescue of the variability phenotype.

Table 1 Detailed statistics used in different figures. It is to be noted that all of the data in figures is from CA1 cells except in figure 4, where it is from both CA3 and CA1 cells.

## Discussion

We have used a mouse model of autism, the fragile X knockout mouse, to investigate cellular correlates of autism in hippocampal CA1 pyramidal neurons. We find that reliability of spiking is impaired in FXS adult animals. We observed an increase in spiking variability both within a cell over multiple trials and between cells in matching trials, in FXS animals. The variability appeared between 6 and 8 weeks of age, consistent with current understanding of autism as a neurodevelopmental disorder. We observed that the increased spiking variability was accompanied by reduced spiking and elevated mAHP currents. We further dissected out the channel contributions to the elevated mAHP and found that SK currents were elevated in adult FXS neurons. However, levels of SK channels were not different between WT and KO mice. To test the hypothesis that elevated SK currents are responsible for unreliable and imprecise spiking in FXS neurons, we used the specific SK blocker apamin and obtained a partial rescue of the phenotype of increased spike variability. In addition to this, we discovered a differential effect of FMRP KO in different hippocampal regions. In agreement with previous published results, we found that there is an increase in excitability in CA3 region of hippocampus, in juvenile animals. At the same age, CA1 cells don’t show any difference in excitability between WT and KO. As the animal becomes older, we observe that the increased excitability seen in CA3 goes away, and for CA1 cells in this age group there is an observation of reduced excitability. This is the first study which has shown that FMRP KO can lead to very different outcomes in different regions of the same brain structure, further highlighting the complexity of the disorder.

### Lowered spike reliability and spike imprecision at cellular intrinsic level in FXS KO hippocampal cells

Our study demonstrates that there is a cell-intrinsic contribution to spiking variability in a mouse model of autism, and implicates increased currents from SK channels as part of the mechanism. This finding takes significance with the failure of mGluR antagonists and blockers to significantly alleviate FXS symptoms in human patients. These findings point to the likelihood that the mGluR theory (Erickson et al., 2017) may be incomplete. Specifically, our finding of within cell variability and between cell variability in FXS cells shows that cells in the mutant animals may form a heterogeneous population of intrinsically variable neurons, the combined effects of which may lead to network-level incoherence.

This variability in neuronal responses over multiple trials may have implications for multiple behaviors. For example, in learning, the stimulus typically needs to be repeated over multiple trials. If the response of a cell at its intrinsic level as well as between cells is variable, the precision of input spiking as well as output activity would be reduced. This would adversely impact learning rules such as STDP and associativity (Markram et al., 1997; Bi and Poo, 1998). Further, elevated SK currents will have a direct impact on the recorded sEPSPs at the soma (Ngo-Anh et al., 2005).

### SK channel currents elevation may underlie previously observed age dependent changes in KO cellular physiology

FXS has many clinical and experimental attributes of a neurodevelopmental syndrome (Nimchinsky et al., 2001; Cruz-Martin et al., 2010; Meredith et al., 2012). An age-dependent decline in learning has been shown in the drosophila Fragile X model (Choi et al., 2010). Theta Burst Stimulation (TBS) LTP has been showed to produced exaggerated LTP in anterior piriform cortex for older KO mice (12-18 months) but not in younger mice (< 6mths)(Larson, 2005). Similar results were seen in Prefrontal Cortex where TBS produced a significant impairment in LTP for 12 month old mice but not in 2 month old animals (Martin et al., 2016). Even in humans it has been shown that the males who have a premutation (have CGG repeats <200) predominantly demonstrated fragile-X associated tremor/ataxia syndrome (FXTAS) among patients who were over 50 years old (Cornish et al., 2008). At the molecular level, an age dependent change has also been shown in protein levels in FXS from P17 mice to adults (Tang et al., 2015)

We observed a similar age dependent change in the spike variability phenotype, which was absent in young KO animals, but manifested in older (>6 week) mice (Figure 3E). Age dependent changes in SK channels have been previously reported such that the levels of SK channels are elevated in older animals as compared to younger ones (Blank et al., 2003). This complements our finding that the AHP difference between WT and KO animals arose in older animals (Fig 5) and that the SK channel underlay the difference in the older animals. There are also evidence of age related changes in Ca^2+^ levels and Ca^2+^ uptake mechanisms, such that basal Ca^2+^ levels are elevated in aging brains (Das and Ghosh, 1996; Kirischuk et al., 1996; Foster and Norris, 1997). Such altered Ca2+ levels would also have a direct impact on the functionality of SK channels. We were not able to directly investigate this possible upstream effect due to the very diverse outcomes of modulating calcium signaling (Figure 8H)

### SK channels may be an FMRP target with multiple physiological and behavioral outcomes

FMRP targets multiple downstream pathways. It not only is a transcription factor but its N-terminus is known to interact with multiple other proteins affecting their functioning (Brown et al., 2010; Deng et al., 2013). Thus mutations in FMRP affect multiple aspects of cellular functioning. Previous RNA-seq results have not shown that SK channels are elevated or affected in FXS (Brown et al., 2001; Darnell et al., 2011). However, the functioning of SK currents might be subject to other indirect effects such as changes in Calcium channel distribution, or in their functioning (Meredith et al., 2007; Ferron et al., 2014). Alterations in SK channels have substantial implications for multiple other phenotypes at the cellular and behavioral level. Previous studies have found that FXS KOs have impaired LTP in CA1 hippocampus (Lauterborn et al., 2007; Lee et al., 2012; Tian et al., 2017). We suggest that this impairment might be partially due to elevated SK currents, which are known to shunt EPSPs and also to increase NMDA Mg^2+^ block (Ngo-Anh et al., 2005). Hence our observation of elevated SK currents is consistent with the reported increase LTP threshold in FXS (Lee et al., 2012). These effects on LTP may have knock-on effects on memory phenotypes in FXS models. For example, a mild impairment in learning due to change in platform location in the Morris water maze test has been found in FXS models (D’hooge et al., 1997; Dobkin et al., 2000).SK currents have been shown to be reduced after learning and this reduction leads decreased variability in spiking events (Sourdet et al., 2003).

### Comparison with previously published studies

Multiple previous studies have found hyper excitability in hippocampus in FXS model mice (Deng and Klyachko, 2016; Luque et al., 2017). We do not find a strong significant difference between KO and WT f-I curves (Fig 3I,3J, 3K and 3L). However, the previously published studies were carried out in the FVB strain of mice and the present study was done on BL/6. Strain dependent differences in FXS have previously been found (Paradee et al., 1999; Dobkin et al., 2000; Lee et al., 2012; Routh et al., 2013). However, (Brager et al., 2012) found a lowering of Input Resistance for KO CA1 cells indicative of reduced excitability. Telias et al 2015 have found similar phenotype as ours in human hippocampal culture cells. In their study, they found that FXS cells were incapable of producing multiple spikes on sustained depolarization at multiple holding potentials, similar to what we observe in this study. It is interesting to note that most excitability analyses and f-I curves use 500 ms current steps, whereas in our study a difference arises after 600 ms, and leads to a reduction in excitability in the KO (Figure 1B and 1I). This suggests that a nuanced interpretation of such excitability studies is desirable, as the responses may have complex temporal features not reducible to a single measure of excitability. Another important point of difference is the age of the animals. Previous published studies have been done on relatively young animals (<P30) as compared to the present study, where adult animals (6-8 wks) have also been used for the experiments. Our own results (Figure 4B and 4C) show that there is a substantial change in spiking variability as well as AHPs as the animal matures. Other studies have also reported an age-dependent effect of expression of plasticity phenotypes in FXS mice (Larson, 2005; Martin et al., 2016). Fragile X Syndrome is known to be a developmental disease where symptoms and many other biochemical and other pathways are known to undergo changes with the age of the animal. Hence it may be important to factor in age-dependence and possibly use a wider range of ages to re-examine some of the conclusions that have been made from FXS models.

Our study is also the first to examine the effect of variability and spiking asynchrony, which again we find to be an age-dependent effect. We postulate that variability in firing of individual cells, which previous studies have not explicitly examined, may be a further complicating factor in analyzing baseline changes arising due to the FMRP mutation.

### FMRP KO has different physiological outcomes in different regions of the hippocampus brain regions

In the present study, we make the important and unexpected finding that FMRP KO has differential cell-physiological effects on different sub sections of the same brain region. In an attempt to replicate previous results, we found that there is hyperexcitability in pyramidal neurons of the CA3 region in juvenile mice (as previously reported by Deng et al 2019) but there was no significant difference in spiking in the CA1 region at the same age. Further, the phenotype reverses for CA1 cells in adult FMRP KO animals such that pyramidal neurons have reduced excitability, whereas for CA3 regions the hyperexcitability goes away. We suggest that these differential effects arise from the balance of effects between the direct effects of FMRP on SK channels, the indirect effects of FMRP on SK via calcium, and the effects of age, again potentially via calcium.

Direct effects of FMRP on SK: Deng et. al. 2019 has shown that absence of FMRP leads to reduced activity of SK channels. They have shown that a physical interaction between FMRP and SK, leads to activation of SK channels. This suggests that FMRP is a direct activator of SK channels. In the case of mutant animals where FMRP is absent, there is a reduction in SK currents leading to hyperexcitability which has been observed both in the present and previous study. (Fig 10A). We see this effect in CA3 cells for KO animals at younger age group.

**Figure 10.**
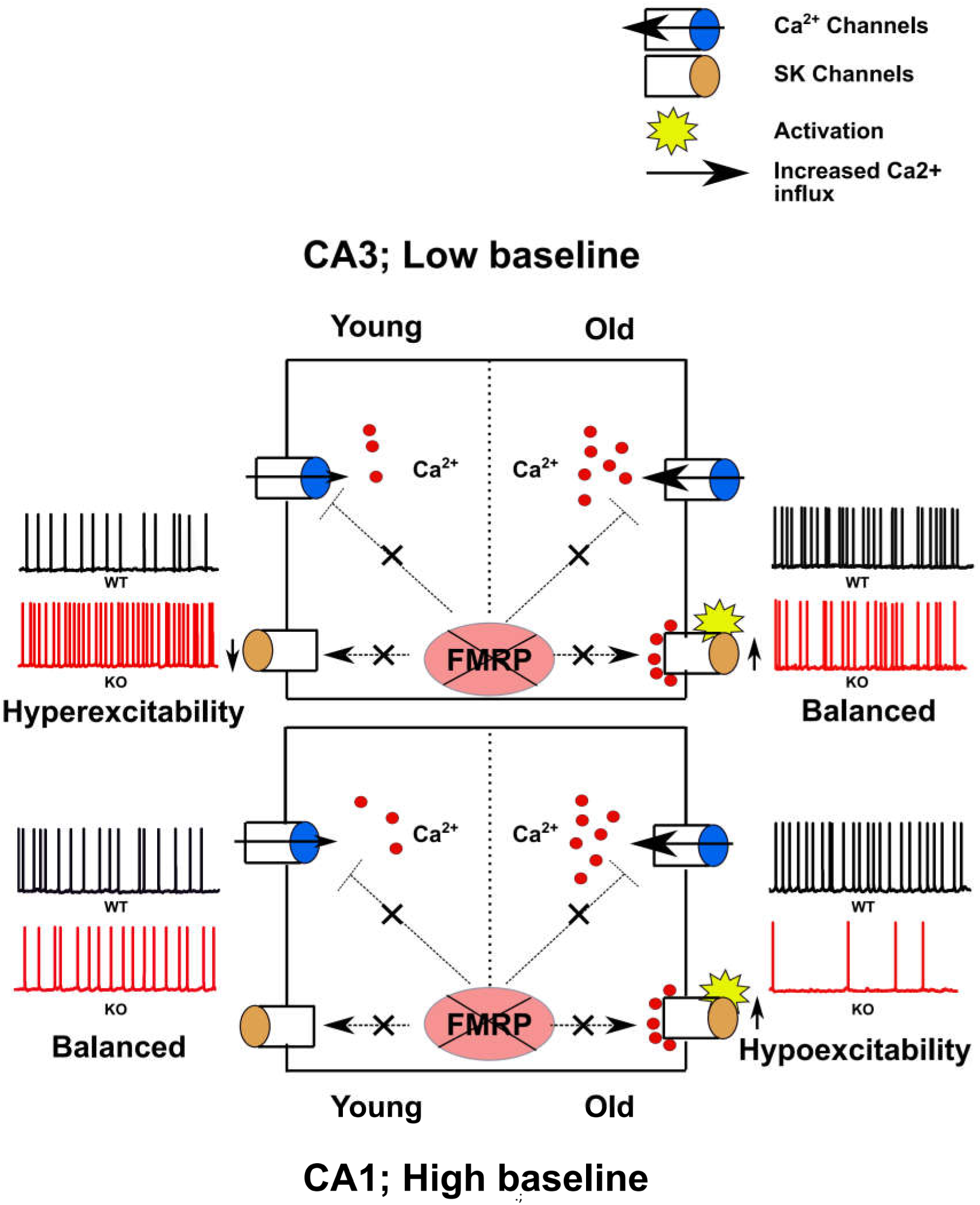
Schematic of a model explaining differential effect of FMRP KO in CA3 and CA1. (A) In young CA3 cells, loss of FMRP leads to reduced activation of SK channels which in turn gives neuronal hyperexcitability. However, as the animal matures there is an increase in calcium influx in these cells. This compensates for this reduced SK activity seen in young CA3 cells leading to a non significant change in excitability for KO cells as compared to WT. (B) In young CA1 cells, both WT and KO cells start with elevated excitability as a result of which no significant differences can be seen between WT and KO. However, as the animal matures, increase in Ca2+ influx brings excitability down for both WT and KO. This decrease in excitability is significantly greater for KO animals as many Ca2+ channels are known to be transcriptionally regulated by FMRP. Under the absence of FMRP, these channels are known to be elevated significantly making KO neurons hypoexcitable as compared to WT.

Indirect effects on SK via calcium: SK channels are dependent on intracellular calcium for their activation. Calcium channels are known targets of FMRP for transcription regulators(Brown et al., 2001; Darnell et al., 2011). Calcium influx via L type Ca2+ channels are known to be elevated in Neural Progenitor Cells (NPCs) (Danesi et al., 2018). Cortical culture neurons have been shown to have significantly elevated basal calcium levels in FXS neuron (Castagnola et al., 2018) and there is increased calcium influx in CA3 cells (Deng et al., 2011, 2013). In reference to human patients, multiple point mutations in Ca^2+^ genes are associated with autism (Contractor et al., 2015; Pinggera et al., 2015; Nguyen et al., 2018). There are multiple other studies in other neuron types which show increased intracellular calcium in autism models(Laumonnier et al., 2006; Krey and Dolmetsch, 2007; Tessier and Broadie, 2011; Pinggera et al., 2015; Limpitikul et al., 2016). These studies are in agreement with our observations in the present study where we see increased SK currents in CA1 cells, which is expected if the intracellular calcium levels are elevated in FXS neurons. (Fig 10A). This increased calcium influx can explain the reduced excitability phenotypes seen in older CA3 and CA1 cells.

Effects of age on calcium: Calcium binding proteins and intracellular calcium dynamics are known to undergo development dependent alterations in FXS neurons (Tessier and Broadie, 2011). Calcium levels and Ca2+ uptake mechanisms, are also known to undergo changes with aging in animals (Das and Ghosh, 1996; Kirischuk et al., 1996; Foster and Norris, 1997). Though there are limited studies available in reference to change in calcium channel levels with age in autism models, we hypothesize that a change in calcium dynamics is consistent with the fact that autism is a neurodevelopment disorder. (Fig 10A and 10B). We see this effect in both CA3 and CA1 regions for KO animal, where in CA3 increased excitability seen in younger animal levels off as the animal becomes adult. On the other hand, because of the increase in calcium, CA1 cells in which excitability was balanced at young age group goes in the direction of hypoexcitability at older age groups.

Based on these findings, we propose a schematic model to explain these differences (Fig 10). In young age group for KO CA3 cells, absence of FMRP leads to hyperexcitability as explained above. As the animal becomes older this effect goes away due to increased calcium influx with age leading to SK activation and balancing out hyperexcitability (Fig 10A). On the other hand for young KO CA1 cells, both WT and KO start with maximum increased excitability where it is not significantly different. As the animal becomes older, increased calcium influx moves KO cells towards hypoexcitability (Fig 10B)

### Defects in single cell physiology may lead to network and population asynchrony

Multiple studies have shown that variability or lack of coordination in spiking exists in FXS at the network level (Arbab et al., 2018; Talbot et al., 2018). In these tetrode-based studies, variability has been shown at the level of cellular firing patterns in the in-vivo network, hence it is difficult to separate cell-intrinsic properties from synaptic and network phenomena. In our study, we use the CV_b_ parameter to gain an understanding about the correlation between cells at a population level. Using this analysis we have tried to estimate the variability which exists between individual neurons responses when actually neurons were recorded one at a time. In contrast to tetrode recordings or field response recordings, where the recorded response is a resultant of multiple neurons and hence is a population response, CV_b_ remains an intrinsic response of neurons depending on its active channels. CV_b_ is an indirect readout of network correlations and should not be inferred as giving actual network correlation information. Clear mechanisms are lacking for this reported asynchrony. However, prior researchers have postulated that synaptic mechanisms may be responsible for the observed asynchrony (Talbot et al., 2018). Our study provides an alternative hypothesis for the mechanism, namely, that the effect is cell-intrinsic and is in part due to altered kinetics of SK channels. Other studies provide evidence in favor of a role for SK channels in mediating precision and synchronous firing. For example, (Deister et al., 2009), showed that SK channels were responsible for mediating precision in autonomous firing in Globus Pallidus neurons. (Schultheiss et al., 2010) showed that dendritic SK channels also control synchronization. Another study by (Kleiman-Weiner et al., 2009) has shown that SK channels play a role in mediating thalamocortical oscillations. A study by (Combe et al., 2018) shows that SK channels contribute to phase locking for spiking of CA1 cells of the hippocampus to slow gamma frequency inputs from Schaffer Collaterals (SC). Thus, alterations in SK channel kinetics in FXS mutants as established by our study might have substantial impacts on phase coding and synchronization of hippocampal cell populations.

### A theory of FXS pathophysiology

An alternate theory to mGLUR theory in explanation of FXS pathophysiology is the ‘discoordination’ theory. The idea behind this theory, is that many of the cognitive impairments seen in FXS are due to altered coordination in spiking of cells in networks, which might be either due to an imbalance in excitation-inhibition in the networks (El Idrissi et al., 2005; Heulens et al., 2012) or due to an intrinsic incoherent tendency of the neurons (Brock et al., 2002; Fenton, 2015; Radwan et al., 2016). Though this theory is preliminary, it may offer a promising alternative point of view from the tranditional mGLUR theory. Some studies have shown that intrinsic changes in cells during FXS effect their E/I balance (Zhang et al., 2014; Lee et al., 2017) and modulation of these channels lead to rescue of FXS phenotype This present study has probed dysfunction of a specific ion channel, SK which was shown to effect the spiking variability at single cell level. It is an addition to the increasing list of ion channels which are known to be affected in FXS and may have an important effect on FXS etiology.

## Acknowledgements

I would like to thank Suranjana Pal, Bishal Basak, Vidya Ramesh, Sarfaraz Nawaz for help with Immunofluorescence experiments. I would also thank Giselle Fernandes, Shreya Das Sharma, Aviral Goel, Sathyaa Subramaniyam, Aanchal Bhatia and Sahil Moza for useful inputs from time to time.

