## Supplementary figures and images for "Impaired reliability and precision of spiking in adults but not juveniles in a mouse model of Fragile X Syndrome"

### Extended data 1-1

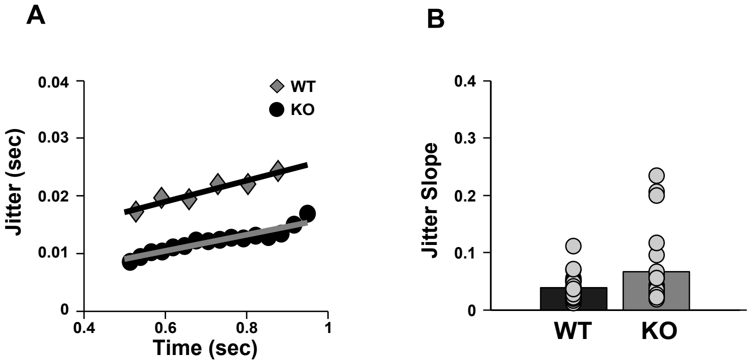

### Extended data 3-1

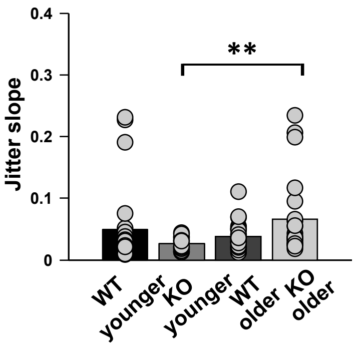

### Extended data 8-1

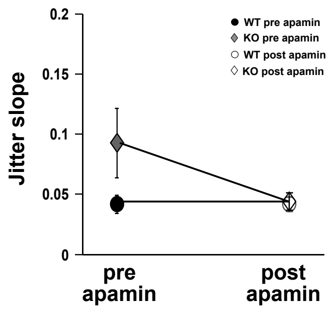
